# Parallel altitudinal clines reveal trends adaptive evolution of genome size in *Zea mays*

**DOI:** 10.1101/134528

**Authors:** Paul Bilinski, Patrice S. Albert, Jeremy J Berg, James A Birchler, Mark Grote, Anne Lorant, Juvenal Quezada, Kelly Swarts, Jinliang Yang, Jeffrey Ross-Ibarra

**Affiliations:** Dept. of Plant Sciences, University of California, Davis, CA, USA; Research Group for Ancient Genomics and Evolution, Dept. of Molecular Biology, Max Planck Institute for Developmental Biology, Tuebingen, DE; Dept. of Biological Sciences, University of Missouri, Columbia, MO, USA; Center for Population Biology, University of California, Davis, CA, USA; Dept. of Evolution and Ecology, University of California, Davis, CA, USA; Dept. of Anthropology, University of California, Davis, CA, USA; Dept. of Horticulture and Agronomy, University of Nebraska, Lincoln, NE, USA; Genome Center, University of California, Davis, CA, USA

## Abstract

While the vast majority of genome size variation in plants is due to differences in repetitive sequence, we know little about how selection acts on repeat content in natural populations. Here we investigate parallel changes in intraspecific genome size and repeat content of domesticated maize (*Zea mays*) landraces and their wild relative teosinte across altitudinal gradients in Mesoamerica and South America. We combine genotyping, low coverage whole-genome sequence data, and flow cytometry to test for evidence of selection on genome size and individual repeat abundance. We find that population structure alone cannot explain the observed variation, implying that clinal patterns of genome size are maintained by natural selection. Our modeling additionally provides evidence of selection on individual heterochromatic knob repeats, likely due to their large individual contribution to genome size. To better understand the phenotypes driving selection on genome size, we conducted a growth chamber experiment using a population of highland teosinte exhibiting extensive variation in genome size. We find weak support for a positive correlation between genome size and cell size, but stronger support for a negative correlation between genome size and the rate of cell production. Reanalyzing published data of cell counts in maize shoot apical meristems, we then identify a negative correlation between cell production rate and flowering time. Together, our data suggest a model in which variation in genome size is driven by natural selection on flowering time across altitudinal clines, connecting intraspecific variation in repetitive sequence to important differences in adaptive phenotypes.

**Author summary:** Genome size in plants can vary by orders of magnitude, but this variation has long been considered to be of little to no functional consequence. Studying three independent adaptations to high altitude in *Zea mays*, we find that genome size experiences parallel pressures from natural selection, causing a linear reduction in genome size with increasing altitude. Though reductions in repetitive content are responsible for the genome size change, we find that only those individual loci contributing most to the variation in genome size are individually targeted by selection. To identify the phenotype influenced by genome size, we study how variation in genome size within a single teosinte population impacts leaf growth and cell division. We find that genome size variation correlates negatively with the rate of cell division, suggesting that individuals with larger genomes require longer to complete a mitotic cycle. Finally, we reanalyze data from maize inbreds to show that faster cell division is correlated with earlier flowering, connecting observed variation in genome size to an important adaptive phenotype.

## Introduction

Genome size varies many orders of magnitude across species, due to both changes in ploidy as well as haploid DNA content [1, 2]. Early hypotheses for this variation proposed that genome size was linked to organismal complexity, as more complex organisms should require a larger number of genes. Empirical analyses, however, revealed instead that most variation in genome size is due to noncoding repetitive sequence and that genic content is relatively constant [3, 4]. While this discovery resolved the lack of correlation between genome size and complexity, we still know relatively little about the makeup of many eukaryote genomes, the impact of genome size on phenotype, or the processes that govern variation in repetitive DNA and genome size among taxa [5].

A number of hypotheses have been offered to explain variation in genome size among taxa. Across deep evolutionary time, genome size appears to correlate with estimates of effective population size, leading to suggestions that drift and ineffective selection permit maladaptive expansion [6] or contraction [7] of genomes across species. A recent evaluation of genome size and the strength of purifying selection among isopods finds evidence supporting this model on a smaller phylogenetic scale [8], but broad-scale phylogenetic analyses fail to find evidence of a correlation between effective population size and genome size, casting doubt on its generality [9, 10]. Other models consider mutation rates, positing that genome sizes evolve to stable equilibria in which the loss of DNA through frequent small deletions is equal to the rate of DNA gain through large insertions. Evidence of the phylogenetic lability of genome size among plants in the family Brassicaceae [11], however, appears inconsistent with this model. Variation in reproductive systems may also explain differences in genome size, as the lower effective population size expected in selfing or asexual species should lead to a reduced ability to purge slightly deleterious novel insertions. Phylogenetic comparisons of repeat abundance and genome size across reproductive systems in *Oenothera*, however, find little support for this hypothesis [12]. In addition to these neutral models, many authors have proposed adaptive explanations for genome size variation. Numerous correlations between genome size and physiologically or ecologically relevant phenotypes have been observed, including nucleus size [13], plant cell size [14], seed size [15], body size [16], and growth rate [17]. Adaptive models of genome size evolution suggest that positive selection drives genome size towards an optimum, due to selection on these or other traits, and that stabilizing selection prevents expansions and contractions away from the optimum [18]. In most of these models, however, the mechanistic link between genome size and phenotype remains unclear [19].

Much of the discussion about genome size variation has focused on variation among species, and intraspecific variation has often been downplayed as the result of experimental artifact [20], or argued to be too small to have much evolutionary relevance [21]. Nonetheless, intraspecific variation in genome size has been documented in hundreds of plant species [21], including multiple examples of large-scale variation [22–24]. Correlations between intraspecific variation in genome size and other phenotypes or environmental factors have also been observed [22, 23, 25], suggesting the possibility that some of the observed variation may be adaptive.

Here we present an analysis of genome size variation in the model system maize (*Zea mays* ssp. *mays*) and its wild relative highland teosinte (*Zea mays* ssp. *mexicana*). Genome size varies as much as 70% between maize and teosinte and genome size within subspecies correlates with both altitude and latitude [23, 26]. Sequencing of the maize reference inbred B73 revealed that the vast majority (85%) of the genome is comprised of transposable elements (TEs) [27], and comparisons between maize and related taxa suggest that variation between species may be explained largely by differences in TE content [28–30]. Within maize, a number of different repeats contribute to variation in genome size. BAC sequencing has identified substantial TE polymorphism among individuals [31, 32], but individuals also vary in the number of auxiliary B chromosomes [33] and large heterochromatic knobs made up of tandem satellite sequences can make up as much as 8% of the genome [34].

We take advantage of parallel altitudinal clines in maize landraces from Mesoamerica and South America to investigate the evolutionary processes and sequence differences underlying genome size variation. Our comparison of flow cytometry data to genotyping reveals evidence that selection has shaped patterns of genome size variation across altitude, and similar analysis of repeat content from low coverage shotgun sequencing identifies an important role for knob variants. We then perform growth chamber experiments to measure the effect of genome size variation on the developmental traits of cell production and leaf elongation in the related wild highland teosinte *Z. mays* ssp. *mexicana*. These experiments find modest support for slower cell production in larger genomes, but weaker support for a correlation between genome size and cell size. Based on these results and reanalysis of published data, we propose a model in which variation in genome size is driven by natural selection on flowering time across altitudinal clines, connecting repetitive sequence variation to important differences in adaptive phenotypes.

## Materials and methods

Unless otherwise specified, raw data and code for all analyses are available on the project Github at https://github.com/paulbilinski/GenomeSizeAnalysis and S1 Table shows the general relationship among samples and analyses; additional details are included below.

### Genome Size

We sampled one seed from each of 77 maize landrace accessions collected across a range of altitudes in Mesoamerica and South America to quantify genome size (S2 Table; [36]). For comparison to maize, we sampled two seeds from 6 and 10 previously collected populations of the wild subspecies *parviglumis* and *mexicana*, respectively (S3 Table; [37]). For our growth chamber experiment, we sampled 201 total seeds from 51 maternal plants collected from 11 populations of *mexicana* (S4 Table and S5 Table). Finally, to assess the error associated with flow cytometry measures of genome size, we used 2 technical replicates of each of 35 maize inbred lines (S6 Table). We germinated seeds and grew plants in standard greenhouse conditions. We collected samples of leaf tissue from each individual and sent material to Plant Cytometry Services (JG Schijndel, NL) for genome size analysis. *Vinca major* was used as an internal standard for flow cytometric measures. Replicated maize lines showed highly repeatable estimates (corr = 0.92), with an average difference of 0.0346pg/1C between estimates.

### Genotyping

We used genotyping-by-sequencing (GBS) [38] data from Takuno *et al.* [36] for maize accessions along altitudinal clines in Mesoamerica and South America. For the 11 *mexicana* populations used in our linear model, we used GBS SNP data from O’Brien *et al.* [39]. All samples were filtered with TASSEL (V5.2.37) [40] to remove sites with *>*40% missing data and individuals with *>*90% missing data, resulting in 170 total individuals with genotyping data for 223,657 sites. We elected to use this per-site cut off as it did not qualitatively change the site frequency spectrum (S1 Fig).

### Kinship and admixture

Kinship matrix calculation was performed using centered identity-by-state (IBS) as implemented in the software TASSEL [41]. We elected to use random imputation in our kinship calculations, as mean imputation biases the estimate of inbreeding within individual [41]. However, we tested both mean and KNN [42] imputation, and our results were robust to both methods. Inbreeding statistics for individual *mexicana* plants were calculated from the diagonal of the randomly imputed kinship matrix.

Admixture analyses were performed using Admixture v1.23 [43]. For admixture analyses we also included additional GBS data from diverse maize inbred lines [44], landraces and teosintes [45] (S2 Table and S4 Table), for a total of 611 individuals before filtering. We filtered individuals and sites as above, but additionally removed one individual (the sample with lowest sequencing depth) of each pair of with an IBS distance closer than 0.07. A Hardy-Weinberg filter was then applied using only outbred genotypes with a read depth between 9-300 using a chi-squared goodness of fit test, p-value *<*0.05. We then thinned sites by linkage disequilibrium, removing lower coverage sites within physical distance less than 1000bp and sites with *r*^2^ >0.8 and significant at p-value <0.05. Only sites with at least 12 high depth genotypes were tested. After filtering, 526 individuals and 18,716 sites remained.

### Shotgun Sequencing

We used whole genome shotgun sequencing to estimate repeat abundance in the same 77 maize landrace accessions and 93 *mexicana* individuals for which we estimated genome size, as well as an additional set of *mexicana* individuals used to validate the approach cytologically (see below, data available on Figshare at DOI 10.6084/m9.figshare.5117827). DNA was isolated from leaf tissue using the DNeasy plant extraction kit (Qiagen) according to the manufacturer’s instructions. Samples were quantified using a Qubit (Life Technologies) and 1ug of DNA was fragmented using a bioruptor (Diagenode) with cycles of 30 seconds on, 30 seconds off. DNA fragments were then repaired with the End-Repair enzyme mix (New England Biolabs), and a deoxyadenosine triphosphate was added at each 3’end with the Klenow fragment (New England Biolabs). Illumina Truseq adapters (Affymetrix) were then added with the Quick ligase kit (New England Biolabs). Between each enzymatic step, DNA was washed with sera-mags speed beads (Fisher Scientific). Samples were multiplexed using Illumina compatible adapters with inline barcodes and sequenced in 3 lanes of a Miseq (UC Davis Genome Center Sequencing Facility) for 150 paired-end base reads with an insert size of approximately 350 bases. The first lane included all maize landraces used for selection studies, the second had the *mexicana* populations used for FISH correlations, and the third included all *mexicana* samples used for analysis of clinal variation.

### Estimating Repeat Abundance

We gathered reference sequences for 180bp knob, TR1 knob, B chromosome, and rDNA repeats from NCBI. CentC repeats were taken from Bilinski *et al.* [46], and chloroplast DNA and mitochondrial DNA were taken from the maize reference genome (v2, www.maizesequence.org). B chromosomes repeats [47] were matched against the maize genome (v2, www.maizesequence.org) using BLAST, and any regions within the repeats that had alignments of greater than 30bp with 80% homology were masked. The remaining unmasked regions with length greater than 70bp were used as a mapping reference for B-repeat abundance. For the transposable element database, we began with the TE database consensus sequences [27, 48]. We then matched sequences against themselves using BLAST and masked any shared regions, retaining unique regions that were at least 70bp in length in our mapping reference. This process was repeated to eliminate regions of secondary homology as well as using knobs and CentC to remove regions of homology to tandem repetitive sequences. We mapped sequence reads to our repeat library using bwa-mem [49] with parameters −B 2 −k 11 −a to store all hit locations with an identity threshold of approximately 80%. We used a minimum seed length of 11 for all repeats as it produced the most reads mapping against the full transposable element database, though our estimates of total repeat abundance (63-71%) are lower than previous estimates [27]. To standardize comparisons of repetitive content across individuals, we first filtered out plastid sequences, then calculated Mb of sequence for each repeat class by multiplying its relative abundance in our sequencing data by genome size converted to base pair values. The correlation between the abundance of each repeat and genome size were as follows: TE = 0.95; 180bp knob = 0.81; TR1 = 0.86. Previous simulations suggest that this estimate has good precision and accuracy in capturing relative differences across individuals [46].

### Repeat Abundance Validation via FISH

We selected two individuals each from 10 previously collected populations of *mexicana* [50] for fluorescence *in situ* hybridization counts of knob content (FISH; S3 Table). FISH probe and procedures closely followed Albert *et al.* [51].

### Clinal Models of Genome Size and Repeat Abundance

We model genome size as a phenotype whose value is a linear function of altitude and kinship (Equation 1). We assume genome size has a narrow sense heritability *h*^2^ = 1, as it is simply the sum of the base pairs inherited from both parents. In our model *P* is our vector of phenotypes, *µ* is a grand mean, *A* is a vector of altitudes included as a fixed effect, *g* represents an additive genetic component modeled as a random effect with covariance structure given by the kinship matrix (**K**), and *ε* captures an uncorrelated error term. The coefficient *β_alt_* of altitude then represents selection along altitude, while the additive genetic (*V_A_*) and error (*V_∊_*) variances are nuisance parameters.

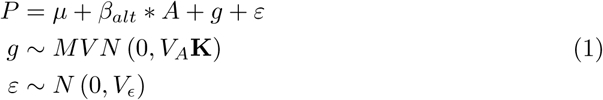

We implemented our linear model in EMMA [52] to test for selection on genome size. In a second model, we then include genome size (GS) as a fixed effect in order to test for correlations between specific repeat classes and altitude conditional on genome size (Equation 2). Estimates of parameters for each model are reported in S8 Table.

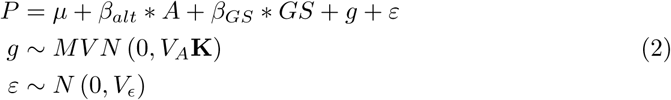

### Growth Chamber Experiment

We conducted a growth chamber experiment to investigate whether genome size variation has an impact on cell production and leaf elongation. We sampled 202 seeds from multiple maternal plants collected in a single high altitude (2408m) *mexicana* population at Tenango Del Aire, chosen because it exhibited the most variation in genome size in our altitudinal transect of *mexicana*. We soaked each fruit in water for 24 hours, then manually removed most of the fruitcase and placed the seed in a Petri dishes inside a growth chamber (23*^°C^*, 16h Light / 8h dark) with cotton balls and water to prevent drying.

Germinated seedlings were transferred to soil pots and into a growth chamber (23*^°C^*, 16h Light / 8h dark). Individually potted seedlings were randomly placed in trays, given fertilized water via bottom watering, and monitored for third adult leaf emergence. We measured leaf length daily for 3 days after the first visible emergence of the third leaf. We clipped the first 8cm of leaf material from the tip of the measured leaf, then extracted a 1cm section which was dipped in propidium iodide (.01mg/ul) for fluorescent imaging (10x magnification, emission laser 600-650, excitation 635 at laser power 6). A minimum of 5 non-overlapping images were taken per leaf sample, horizontally across the leaf segment if possible. Cell length was measured for multiple features, including stomatal aperture size and rows adjacent to stomata. Lengths across different features were highly correlated, so stomatal aperture size was used as the repeated measure of cell lengths in the growth model.

### Modeling the Effect of Genome Size on Cell Production

We model leaf elongation rate (LER) as the product of cell size (CS) and the rate of cell production (CP):

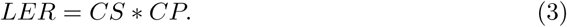

The multiplicative expression in Equation 3 is linearized by taking the natural logarithm on both sides of the equation, and model-fitting is performed on the log scale. We hypothesize that genome size affects LER only through its effects on CS and CP. The strategy for estimation of genome-size effects is illustrated by path diagrams shown in S2 Fig, where additional details are given. We adopt a computational Bayesian approach for parameter estimation, incorporating seedling and maternal random effects in models that make use of the hierarchical dataset structure (cells and days of growth within seedlings, seedlings within maternal parents). The signs and magnitudes of our estimated effects, and therefore our conclusions, are sensitive to different specifications of prior information. We identified previous averages for maize stomatal cell size and daily leaf elongation rate (CS=0.003 cm, LER=4.0-4.8cm/day or 2mm/hr) [53–56], and incorporated these into informative priors for the random effects. Because our model shows prior sensitivity, we also identify prior means for which the sign of the relationship between genome size, cell production rate (*β_GS_*), and cell size (*γ_GS_*) changes (S3 Fig). We generated posterior samples of model parameters using JAGS, a general-purpose Gibbs sampler invoked from the R statistical language using the library rjags [57]. We allowed for a burn in of 200,000 iterations and recorded 1,000 posterior estimates by thinning 500,000 iterations at an interval of 500.

### Analysis of Maize SAM Cell Number and Flowering Time

To evaluate evidence for a relationship between cell production rate and flowering time, we used flowering time and meristem cell number data for 14 maize inbred lines from Leiboff *et al.* [58]. Because meristems were sampled at an identical growth stage and time point, differences in cell number should reflect differences in the rate of cell production. We fitted a mixed linear model to estimate the best linear unbiased estimates (BLUEs) of the cell counts for each growth period separately:

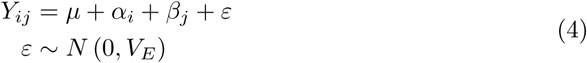

In this model, *Y_ij_* is the cell count value of the *i^th^* genotype evaluated in the *j^th^* replicate; *µ*, the overall mean; *α_i_*, the fixed effect of the *i^th^* genotype; *β_j_*, the random effect of the *j^th^* block; and *ε*, the model residuals.

Each line’s genotype at trait-associated SNPs for the candidate genes BAK1 and SDA1 [58] was considered as a fixed effect and replication as a random effect. We then fitted mixed linear models to study the relationship of flowering time and cell counts by controlling for population structure and known trait-associated SNPs:

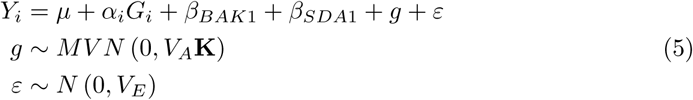

Here *Y_i_* is the flowering time (days to anthesis) of the *i^th^* genotype; *μ*, the overall mean; *α_i_*, the fixed effect of the *i^th^* Genotype; *β_BAK1_* and *β_SDA1_* the fixed effects of the BAK1 and SDA1 loci; *g* a random effect modeled with a covariance structure given by the kinship matrix **K**; and *ε* an uncorrelated error. The additive genetic (*V_A_*) and environmental (*V_E_*) variances are nuisance parameters.

Cell counts were included as fixed effects and the standardized genetic relatedness matrix was fitted as a random effect to control for the population structure [59]. The genetic relatedness matrix was calculated using GEMMA [60] from publicly available GBS genotyping for these lines (AllZeaGBSv2.7 at www.panzea.org, [40]). In the calculation, we used 349,167 biallelic SNPs after removing SNPs with minor allele frequency <0.01 and missing rate >0.6 using PLINK [61].

## Results

We sampled 77 diverse maize landraces from across a range of altitudes in Meso- and South America (S2 Table). Flow cytometry of these samples revealed a negative correlation with altitude on both continents (Fig. 1A, r=-0.51 and -0.8, respectively, p-value <0.001). We used low-coverage whole-genome sequencing mapped to reference repeat libraries to estimate the abundance of repetitive sequences in each individual with estimated genome size, and validated this approach by comparing sequence-based estimates of heterochromatic knob abundance to fluorescence *in situ* hybridization (FISH) data from *mexicana* populations (Fig. 2 and S4 Fig; see Methods for details). Consistent with previous work [28, 35, 62], transposable elements and heterochromatic knobs contributed most to variation in genome size across our maize samples (Fig. 1) and we find only a weak positive correlation between B chromosome abundance and altitude (p-value *>* 0.05) (S6 Fig). We observed substantial variation among landraces in the abundance of individual transposable element families (S5 Fig), and both transposable elements as a whole and heterochromatic knobs showed clear decreases in abundance with increasing altitude in both Meso- and South America (TE r=-0.57, -0.72; 180bp knob r=-0.48, −0.83; TR1 knob r=-0.66, -0.81; p-value < 0.001), mirroring the pattern seen for overall genome size (Fig. 1).

**Fig 1.**
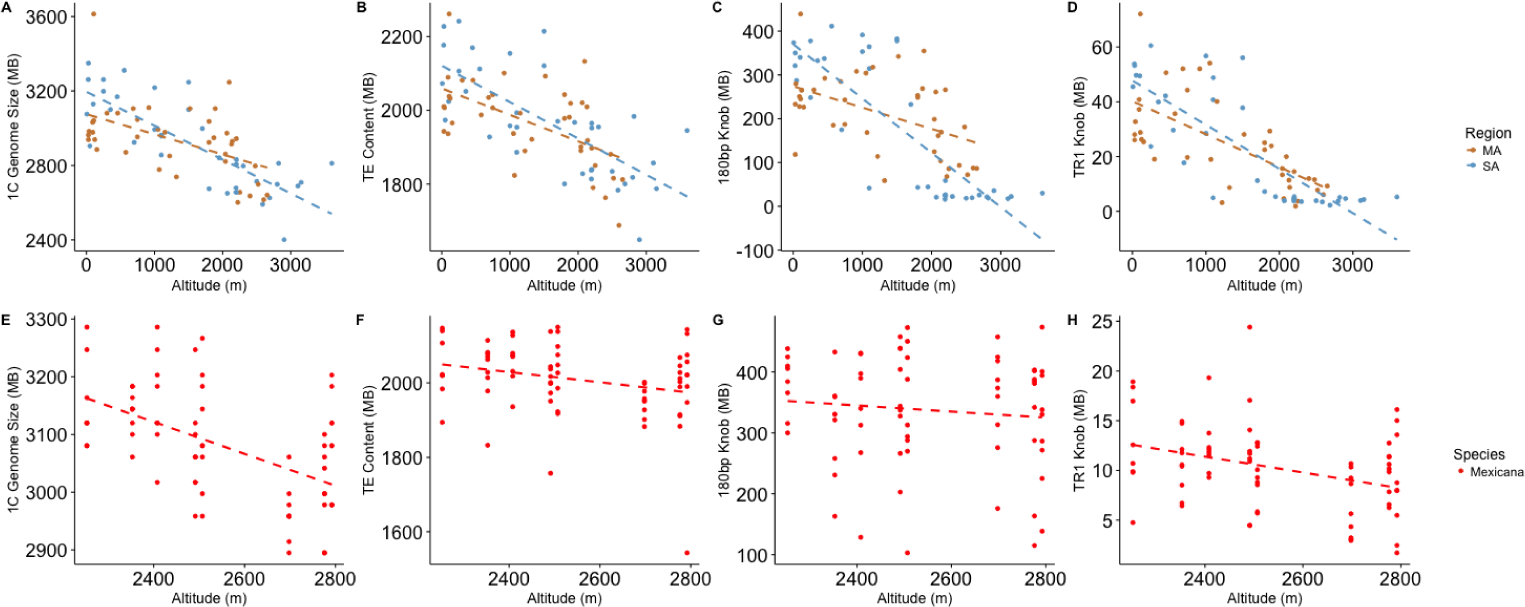
Genome size and repeat content by altitude in *Zea* taxa. (A-D) Maize landraces from Mesoamerica (MA) or South America (SA). (E-H) Highland teosinte *Z. mays* ssp. *mexicana*. Only teosinte populations above 2000m that do not show admixture (see text) are included. (A,E) total genome size, (B,F) total transposable element content, (C,G) 180bp knob repeat content, (D,H) TR1 knob repeat content. Dashed lines represent the best fit linear regression.

**Fig 2.**
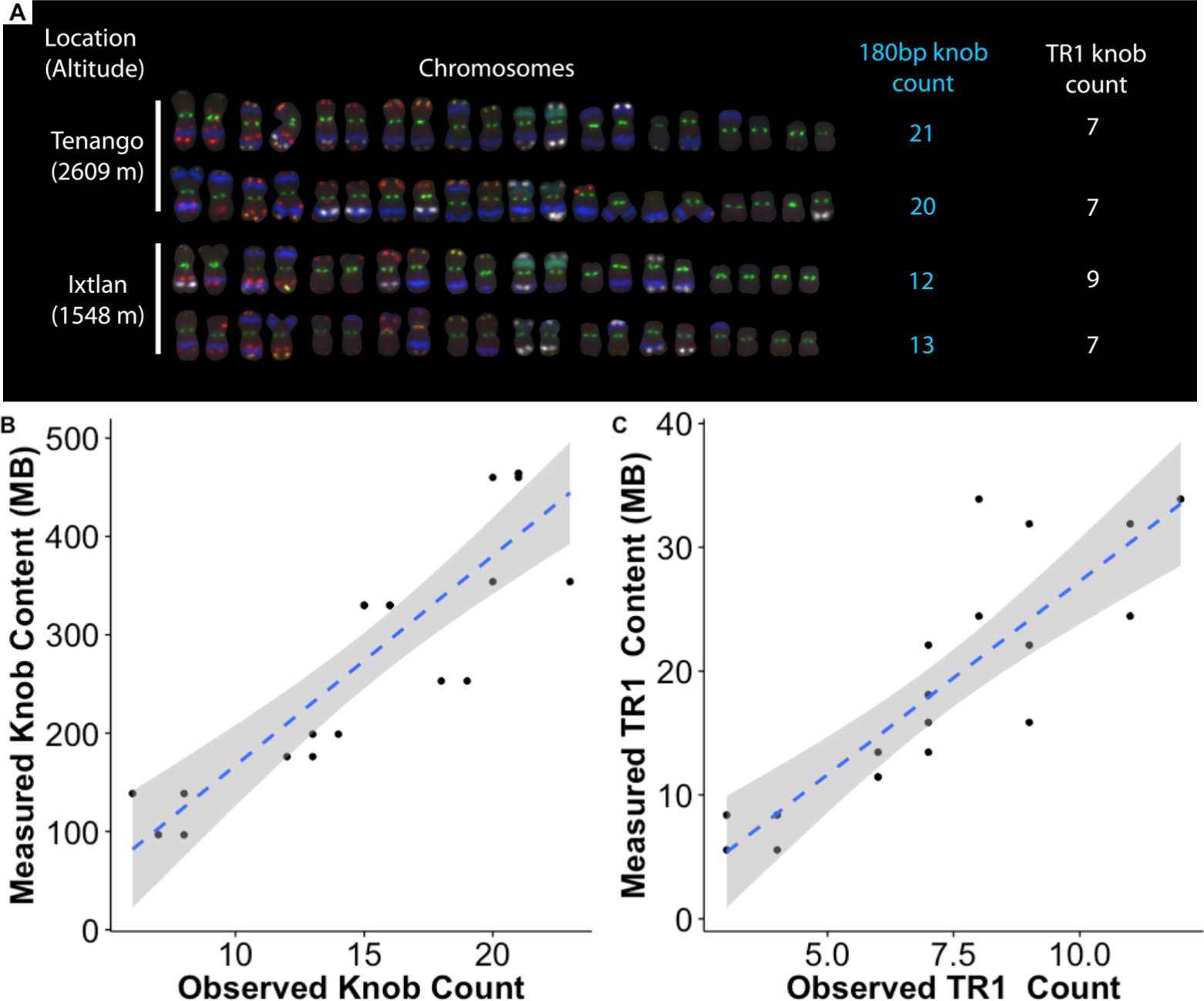
Knob content in highland teosinte estimated using FISH and low-coverage sequencing. (A) FISH from four *Z. mays* ssp. *mexicana* individuals, sampled from the highest and lowest altitude populations. Counts of cytological 180bp (blue) and TR1 (white) knobs are shown to the right of each individual. Other stained repeats are CentC and subtelomere 4-12-1 (green), 5S ribosomal gene (yellow), Cent4 (orange), NOR (blue-green), and TAG microsatellite 1-26-2 and subtelomere 1.1 (red). For further staining information, see [51]. (B) Plot of the population-level correlation between 180bp knob counts and sequence abundance for 20 *mexicana* individuals. 180bp knob r = 0.88, TR1 knob r = 0.86

We next sought to evaluate whether the observed clines in genome size and repeat abundance simply reflected underlying genetic differences due to population structure, or could be better explained by natural selection along an altitudinal cline. We adopted an approach similar to Berg and Coop [63], modeling genome size as a quantitative trait that is a linear function of relatedness and altitude (see Methods, Equation 1). Across maize landraces, we rejected a neutral model in which genome size is unrelated to altitude, estimating a decrease of 108Kb and 154Kb in mean genome size per meter gain of altitude in Meso- and South America, respectively (S8 Table). We then evaluated whether selection has acted on individual repeats, treating abundance of each repeat class as a quantitative trait in a comparable model that includes genome size as a covariate (Methods, Equation 2). In both Meso- and South America, TR1 knobs showed evidence of selection, while 180bp knob also showed evidence of selection in South American landrace germplasm (S8 Table). Finally, our models for total transposable element content in both Mesoamerican and South American maize were just above the 0.05 significance threshold, and the number of individual TE families showing significant correlations with altitude was no greater than expected by chance (46/1156, binomial test p-value >0.05).

The wild ancestor of maize, *Zea mays* ssp. *parviglumis* (hereafter *parviglumis*), grows on the lower slopes of the Sierra Madre in Mexico. A related wild teosinte, *Zea mays* ssp. *mexicana* (herafter, *mexicana*), diverged from *parviglumis* ≈60,000 years ago [64] and has adapted to the higher altitudes of the Mexican central plateau [65]. We sampled leaves and measured genome size of two individuals each from previously collected populations of both subspecies (6 *parviglumis* populations and 10 *mexicana* populations) [37, 50]. Though both subspecies exhibit considerable variation, *mexicana* had a smaller average genome size than *parviglumis* (S7 Fig; one tailed t-test p-value <0.05), consistent with our observations of decreasing genome size along altitudinal clines in Mesoamerican and South American maize.

To evaluate clinal patterns across populations of highland teosinte in more detail, we sampled multiple individuals from each of an additional 11 populations of *mexicana* across its altitudinal range in Mexico (S4 Table). Genome size variation across these populations revealed no clear relationship with altitude (S8 Fig), but genotyping data [39] revealed consistent evidence of genetic separation (S9 Fig) and higher inbreeding coefficients (two-sided t-test p-value <0.001) in the three lowest altitude populations (see Methods). These three populations are also phenotypically distinct and relatively isolated from the rest of the distribution (A. O’Brien pers. communication). We thus excluded these three populations, applying our linear model of altitude and relatedness to 70 individuals from the remaining 8 populations. After doing so, we find a negative relationship between genome size and altitude in *mexicana* (Fig. 1E, p-value <0.001) of similar magnitude to that seen in maize (loss of 270Kb/m), suggesting parallel patterns of selection across *Zea*. In agreement with our results in maize, TR1 knob repeats showed evidence of selection after controlling for their contribution to genome size (S8 Table), though 180bp knob repeats did not. We found no evidence for selection on TE abundance after controlling for genome size, and none of the sequence from *mexicana* mapped to our B-repeat library.

When grown in a common environment, both highland maize and highland teosinte flower earlier than their lowland counterparts [66, 67], and previous work has shown that selection for early flowering time in maize results in a concomitant reduction in genome size [68]. Extrapolating from this observation, we reasoned that genome size might be related to flowering time through its potential effect on the rate of cell production and consequently development. To test this hypothesis, we performed a growth chamber experiment to measure leaf elongation rate, cell size, and genome size using 201 *mexicana* individuals from 51 maternal families sampled from a single natural population (see Methods). Individual plants varied by as much as 1.13Gb in 2C genome size, with observed leaf elongation rate (LER) varying from 1 to 8 cm/day (mean 4.56cm/day; S9 Table). We designed a Bayesian model of leaf elongation as a function of cell size, cell production rate, and genome size (see Methods). Our posterior parameter estimates suggest a weak but positive relationship between genome size and cell size (*γ_GS_*; Fig. 3A) and a negative relationship between genome size and cell production rate (*β_GS_*; Fig. 3B). We found that our inferences were sensitive to prior specifications for leaf elongation rate and cell size (S3 Fig), but prior means ≥ 4cm/day for leaf elongation rate combined with prior means ≤ 0.003cm for CS, returned reliably negative relationships between genome size and cell production rate (see Methods).

**Fig 3.**
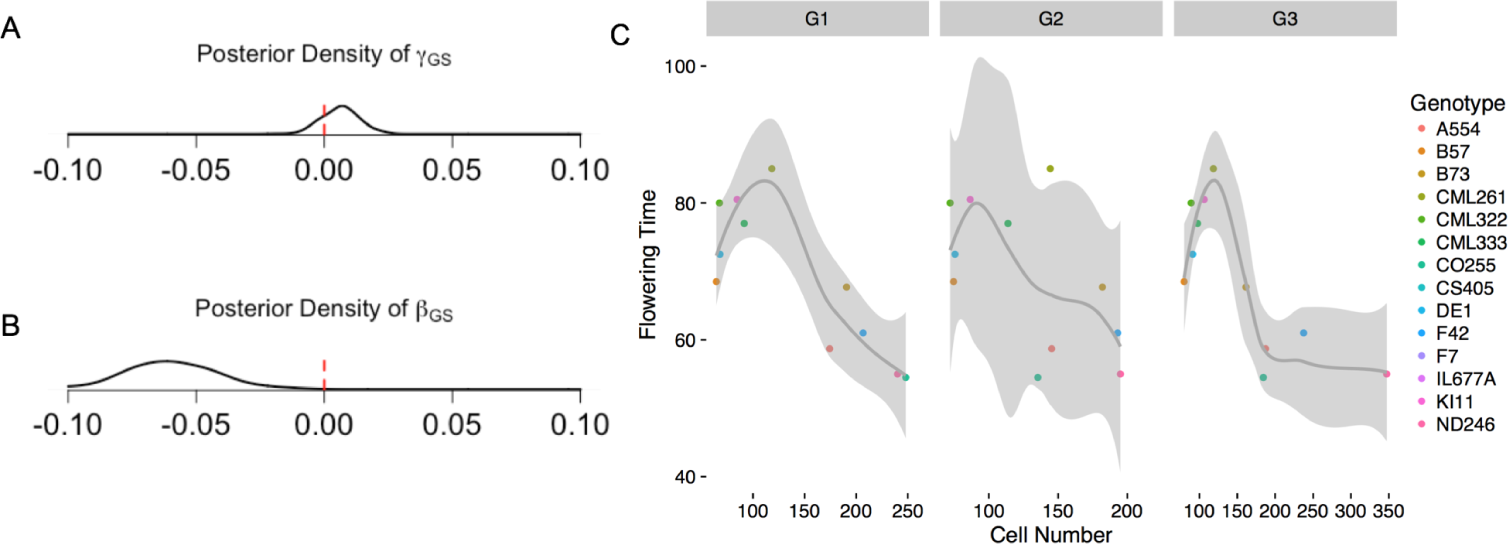
(A,B) Posterior densities of effects of genome size on cell size and cell production rate (*γ_GS_* and *β_GS_*, respectively) from a model with prior mean stomatal cell size of 30 microns and leaf elongation rate of 4cm/day. (C) Smoothed splines showing the relationship between flowering time and SAM cell number in inbred maize accessions. Measurements for cell number are shown for each of three growth phases (G1, G2, G3). Data from Leiboff *et al.* [58].

Recent work exploring shoot apical meristem (SAM) phenotypes across 14 maize inbred lines [58] allowed further exploration of our hypothesized connection between cell production and flowering time. Because Leiboff *et al.* sampled SAM at equivalent growth stages, we interpreted variation in cell number as representative of differences in cell production rate among lines. We re-analyzed these data to investigate whether the cell number reported in each SAM was correlated with flowering time (Fig. 3C). After estimating genetic values for each inbred line used and correcting for population structure and the effects of two candidate genes (see Methods), we find a negative correlation between flowering time and cell production across all three developmental stages sampled (slopes of −0.11, −0.08, and −0.08 and p-values < 0.01, <0.001, and 0.170, respectively). Such negative correlations suggest that a higher number of cells in the seedling SAM — and thus a faster rate of cell production — may lead to earlier flowering time.

## Discussion

### Genome size and repeat abundance

We report evidence of a negative correlation between genome size and altitude across clines in Meso- and South America in both maize and its wild relative highland teosinte (Fig. 1). Genetic evidence suggests that maize colonization of highland environments was independent in Mesoamerica and South America [69], and while the populations share a number of adaptive phenotypes, they exhibit little evidence of convergent evolution at individual loci [36]. The teosinte subspecies *mexicana* is also found in the highlands of Mesoamerica [65], likely after its split from the lowland teosinte *parviglumis* ≈60,000 years ago, long before maize domestication [64]. Previous investigations of genome size have also identified altitudinal clines in maize and teosinte [23, 26] (but see Rayburn *et al.* [70] for a counterexample in the U.S. Southwest), suggesting that this observation is general and not an artifact of our sampling. Although we find altitudinal trends in genome size across all three clines, our initial evaluation of genome size in highland teosinte found no significant correlation with altitude, due primarily to the small genomes observed in the three lowest altitude populations (S8 Fig). We excluded these three *mexicana* populations because they showed higher levels of inbreeding than other *mexicana* populations as well as evidence of shared ancestry with *parviglumis* (S9 Fig). These populations are nonetheless interesting and worthy of future investigation, as their genome size is smaller than either *parviglumis* or high altitude *mexicana*, but their knob content does not differ from other *mexicana* populations, suggesting perhaps that inbreeding or admixture may have affected transposable element or other repeat abundance.

Our results suggest the best explanation for the observed clines in genome size is natural selection. Several authors have identified ecological correlates of variation in plant genome size and argued for adaptive explanations of such clines [23, 26, 71], but did not correct for relatedness among individuals or populations. We employ a modeling approach that considers genome size a quantiative trait and uses SNP data to generate a null expectation of variation among populations, allowing us to rule out stochastic processes and instead pointing to the action of selection in patterning clinal differences in genome size. Alternative explanations for our observations, including mutational biases and TE expansion, are unlikely. Plants grown at high altitudes are exposed to increased UV radiation and UV-mediated DNA damage may lead to higher rates of small deletions [72]. But because UV damage causes small DNA deletions, it is unlikely to generate the gigabase-scale difference we see across altitudinal clines in the short time since maize arrived in the highlands [73]. And while expansion or replication of TE in lowland populations could lead to increased rates of insertion and larger genome size, our analysis of reads mapping to individual TE families finds no evidence that this has occurred in a widespread manner. Moreover, genome size estimates from the direct wild ancestor of domesticated maize (the lowland teosinte *parviglumis*) suggest that smaller highland genomes are the derived state.

Having concluded that natural selection is the most plausible explanation for decreasing genome size at higher altitudes, we then asked whether these observations were the result of selection on genome size itself or merely a consequence of selection on specific repeat classes. We find no evidence of selection on B repeats, consistent with relatively mixed signals found in previous literature [62]. We also find little evidence of selection on TEs after controlling for genome size. Because individual TEs are relatively small, however, models of polygenic adaptation lead us to expect that such loci are unlikely to show a strong signal [74]. Nonetheless, TEs show the strongest overall correlation with genome size, suggesting that frequency of small deletions of individual elements are likely a major contributor to genome size change across populations. In contrast to TEs, in both maize and teosinte the 350bp TR1 knob repeat shows greater differentiation in abundance across altitude than can be explained by population structure alone, even after accounting for changes in total genome size. The 180bp knob shows a similar strong decline in abundance in maize landraces, but is only statistically significant in the analysis of landraces in South America. Selection on genome size might be expected to act especially strongly on knobs because each locus contains many megabases of repeats and thus represents a large contribution to variation in genome size. In contrast, individual transposable elements are thousands of times smaller than heterochromatic knobs, and selection on such small-effect variants is not expected to show a signal much different from the overall phenotype. These results are surprising, however, given the selfish nature of knobs and their ability to distort segregation ratios in female meiosis in the presence of a driving element known as abnormal chromosome 10 (Ab10) [75]. While our genotyping data do not include markers diagnostic of Ab10, previous analyses show that selection along altitudinal gradients has been sufficient to decrease the frequency of at least one allele of the drive locus itself [76]. It is not entirely clear why we see more evidence of selection on the TR1 knob variant, which contributes nearly an order of magnitude fewer base pairs to the genome. The TR1 variant generally shows weaker drive, but has been shown to compete successfully against the 180bp variant [77]. It is thus possible that the weaker drive of TR1 makes it more susceptible to selection on overall genome size, and that the subsequent decrease in TR1 abundance may increase drive of the 180bp knob variant, potentially ameliorating the effects of selection against 180bp knobs at higher altitude. Finally, while we see decreasing abundance of both knob variants with increasing altitude, we note that knobs alone are not driving the overall signal: rerunning our model for genome size after removing base pairs attributable to both knob repeats still finds evidence of selection on genome size in all three clines (Mesoamerica p-value= 0.029; South America p-value= 0.04; *Mexicana* p-value = 0.02).

### Genome size and development rate

Several authors have hypothesized that genome size could be related to rates of cell production and thus developmental timing [75, 79]. We tested this hypothesis in a growth chamber experiment in which we measured leaf elongation rates across individuals from a single population of highland teosinte that exhibited wide variation in genome size. Our approach to characterizing the effect of genome size on the rate of cell production is consistent with scaling laws proposed in a recent study of the relationships between genome size, cell size, and cell production rate [80] (see Methods). We found only weak evidence for a positive correlation between genome size and cell size, a result that contrasts with the findings of many authors who have reported more definitive positive correlations between genome size and cell size across species [81, 82]. One potential explanation for this result may be found in recent work in *Drosophila* where larger repeat arrays were shown to lead to more compact heterochromatin despite the physical presence of more DNA [83]. We speculate that such an effect may ameliorate some of the physical increase in chromosome size due to the expansion of certain repeats, especially tandem arrays such as those found in dense heterochromatic knobs.

In support of the hypothesis that smaller genomes may enable more rapid development, our leaf elongation model indicates a negative correlation between genome size and cell production rate in our highland teosinte population. Though these results showed strong prior sensitivity, the sign of the relationship between genome size and cell production rate did not change for prior mean values of leaf elongation rate within the range of those published for maize (from 4.6 cm/day [56] to 12 cm/day [84]), all equal to or larger than the rates observed in our experiment. Additional evidence comes from a recent study by Tenaillon *et al* [85], who also find a negative correlation between the rate of leaf elongation and genome size, albeit one that does not survive statistical correction for population structure.

We hypothesize that selection on flowering time is the driving force behind our observed differences in genome size. Larger genomes require more time to replicate [71], and slower rates of cell production in turn may lead to slower overall development or longer generation times [79]. Slower cell production is unlikely to be directly limiting to the cells that eventually become the inflorescence, as only relatively few cell divisions are required [86]. However, signals for flowering derive from plant leaves [87, 88], and slower cell production will result in a longer time until full maturity of all the organs necessary for the plant to flower. Highland populations of both maize and teosinte flower earlier than lowland populations [66, 89], and experimental work in maize reveals a genetic correlation of ≈0.14 between flowering time and genome size (data from Rayburn *et al.* [68] assuming heritabilities of *h*^2^ = 0.8 for flowering time and *h*^2^ = 1 for genome size). Consistent with this hypothesis, maize plants with more cells in their SAM at a given developmental stage (and thus faster rates of cell production) appear to also exhibit earlier flowering [58]. Future efforts to experimentally connect genome size to both cell production and flowering time within a single panel will be important to definitively establish a mechanistic connection between genome size and flowering time. In addition to flowering time, the metabolic requirements of nucleotide synthesis could play a selective role in determining plant genome size variation. Nucleotide synthesis requires substantial nitrogen and phosphorous, and it has been argued that selection for rapid growth in nutrient-poor environments may act to reduce genome size [90]. Indeed, phylogenetic comparisons find a significant correlation between nitrogen content (but not phosphorous) and genome size among *Primulina* growing in nutrient-limited karst soils [25]. We are unaware, however, of any meaningful correlations between nitrogen or phosphorus concentration and altitude across either our Mesoamerican or South American clines, suggesting that soil nutrients are unlikely to completely explain the patterns we observe.

## Conclusion

The causes of genome size variation have been debated for decades, but these discussions have often disregarded adaptive explanations and ignored intraspecific variation. Our results suggest that differences in optimal flowering times across altitudes are likely indirectly effecting clines in genome size due to a mechanistic relationship between genome size and cell production and developmental rate. We also show that selection on genome size has driven changes in repeat abundance across the genome, including significant reductions in individual repeats such as knobs that contribute substantially to variation in genome size. We speculate that our observations on genome size and cell production may apply broadly across plant taxa. Intraspecific variation in genome size appears a common feature of many plant species, as is the need to adapt to a range of abiotic environments. Cell production is a fundamental process that retains similar characteristics across plants, and genome size is likely to impact cell production due to the limitations in replication kinetics that result from having a larger genome. Together, these considerations suggest that genome size itself may be a more important adaptive trait than has been previously believed, and that the phenotypic effects of genome size may have consequences for the evolution of individual repeats.

## Supplementary Information

**S1 Fig.**
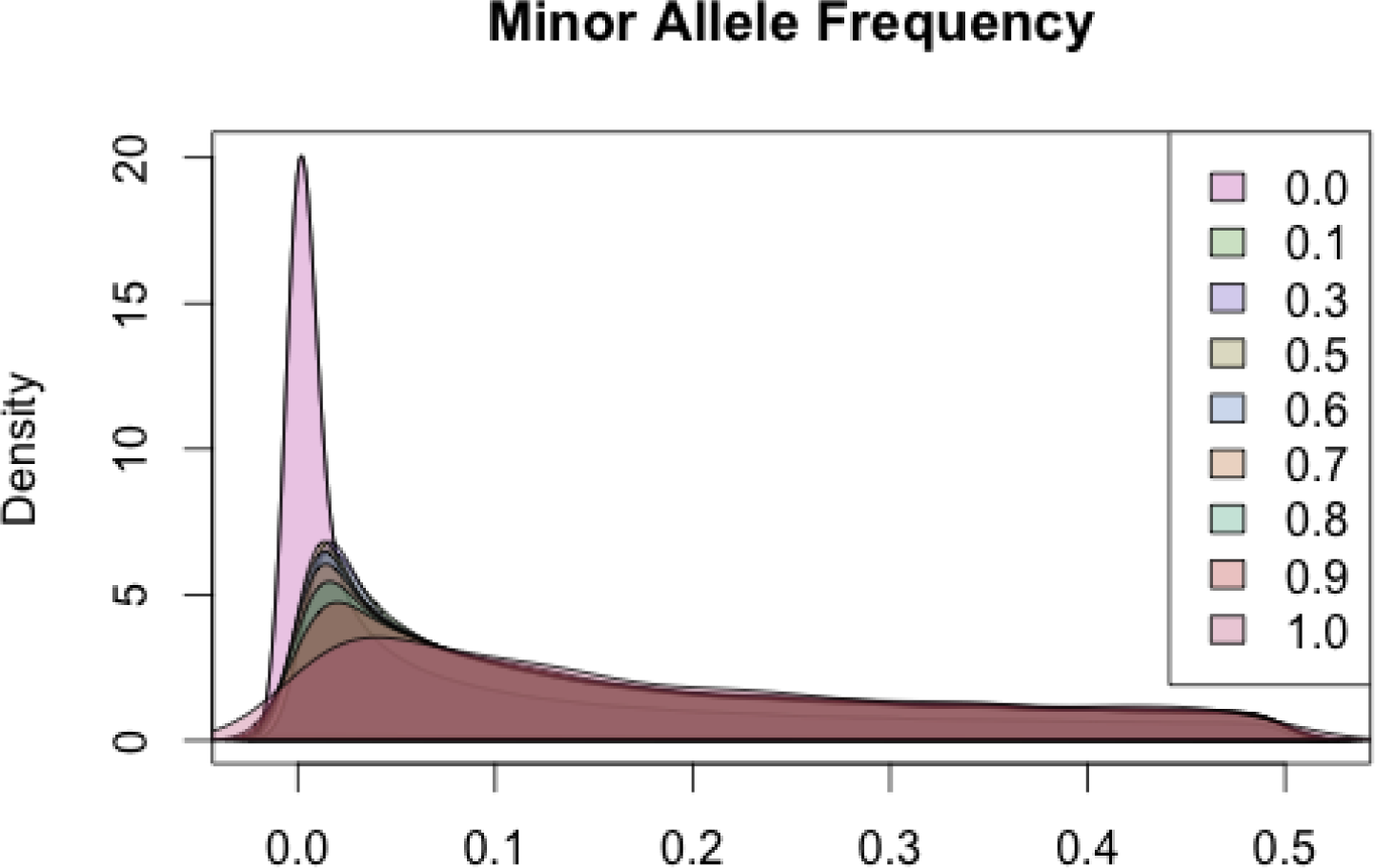
Plot of reads mapping to B chromosome specific repeats in maize landraces. Key indicates percent of site coverage, ranging from unfiltered (0.0) to a requirement of full data presence across individuals (1.0). The spectrum begins to shift after a 60% site coverage requirement.

**S2 Fig.**
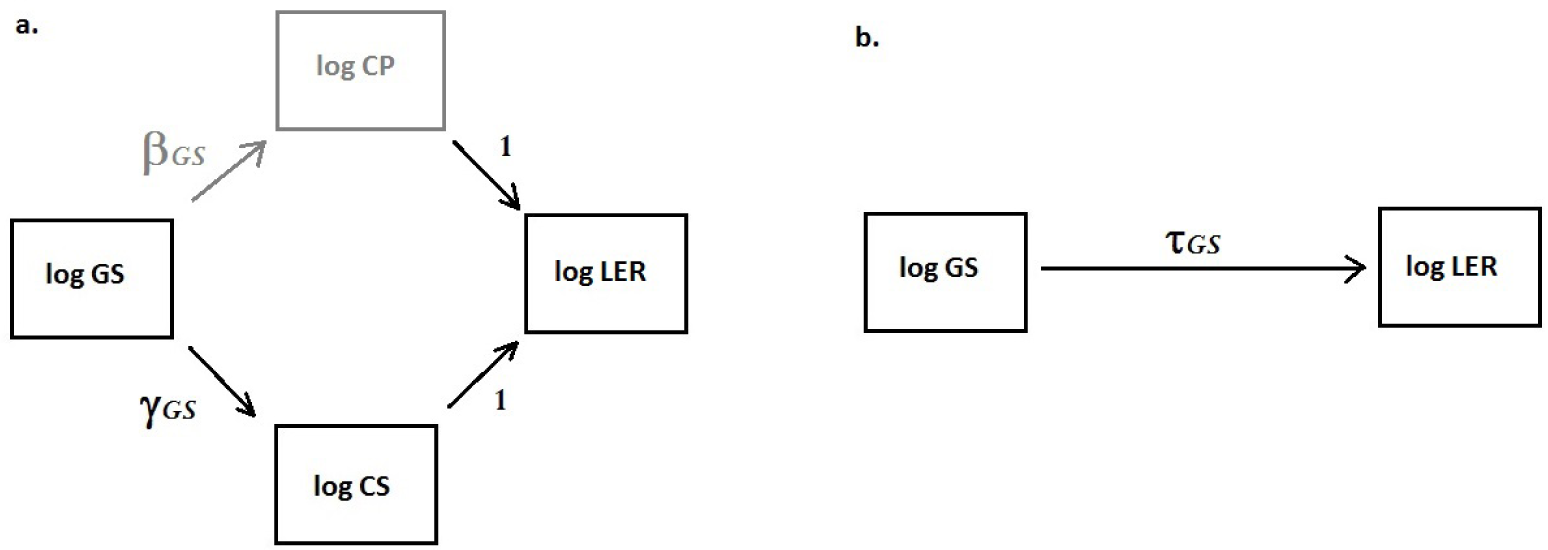
Path models for estimation of genome size effects. Arrows indicate predictor-outcome relationships and are annotated with model coefficients (slopes) from equations 9-11. (A) Genome Size (GS) predicts Leaf Elongation Rate (LER) through the mediators Cell Size (CS) and Cell Production rate (CP). CP, shown in grey, is not directly observed. The unit coefficients connecting log LER with log CP and log CS reflect the assumption LER = CS * CP (equation 3). (B) Marginal model for the effect of GS on LER (equation 10).

**S3 Fig.**
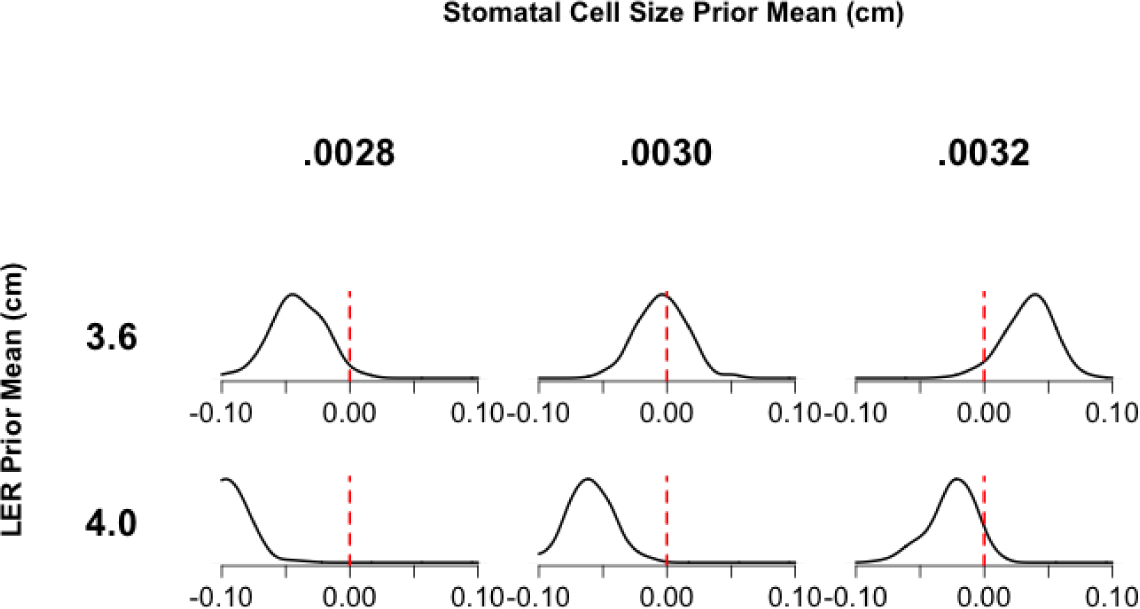
Effect of LER and Stomatal Cell Size Priors on Posterior Density of the Cell Production Coefficient *β_GS_*.

**S4 Fig.**
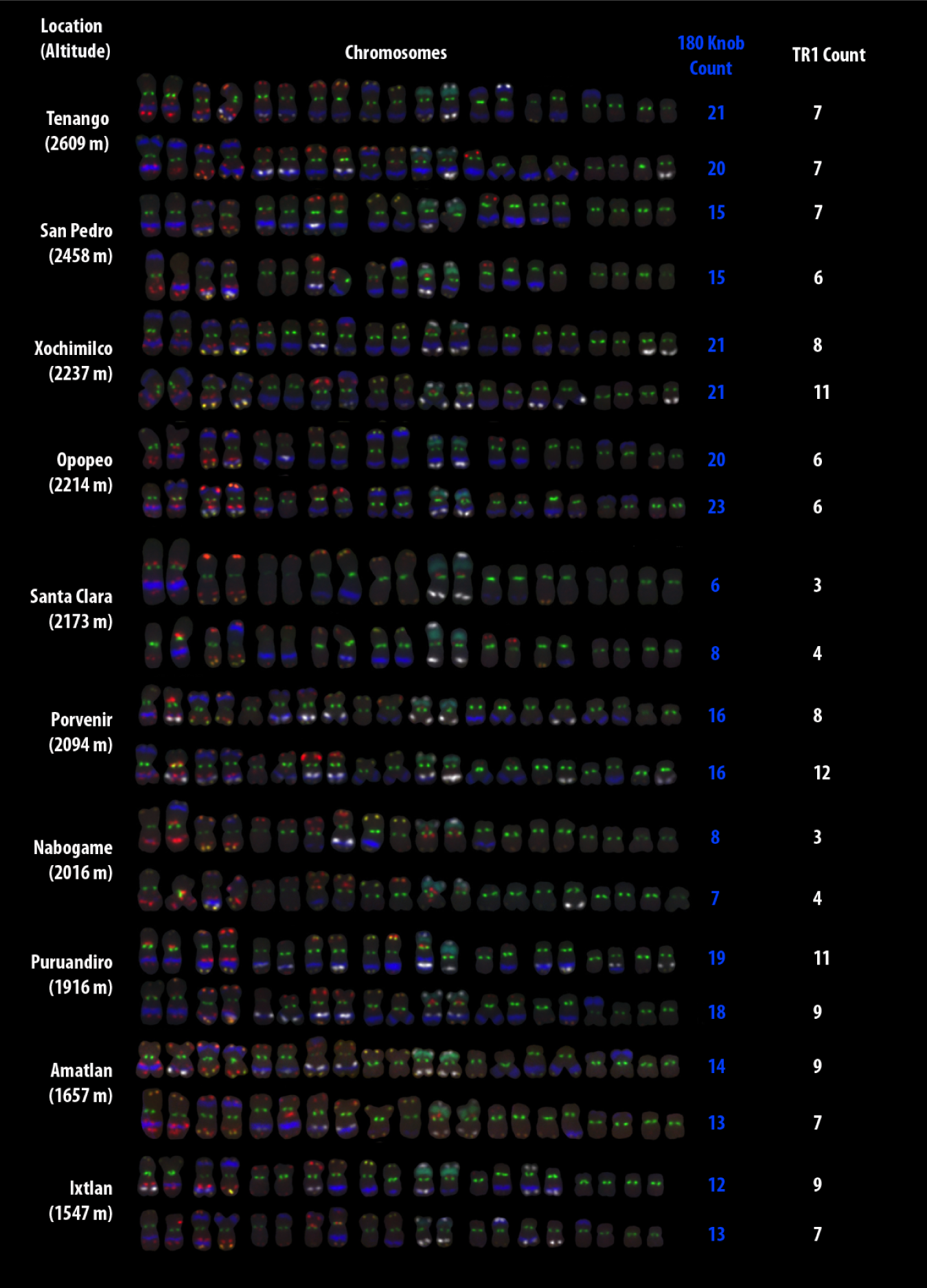
Knob content in highland teosinte estimated using FISH and low-coverage sequencing, showing all sampled individuals referenced in Fig. 2. Counts of cytological 180bp knobs (blue) and TR1 knobs (white) are shown to the right of each individual. Other stained repeats are CentC and subtelomere 4-12-1 (green), 5S ribosomal gene (yellow), Cent4 (orange), NOR (blue-green), and TAG microsatellite 1-26-2 and subtelomere 1.1 (red). For further staining information, see [51].

**S5 Fig.**
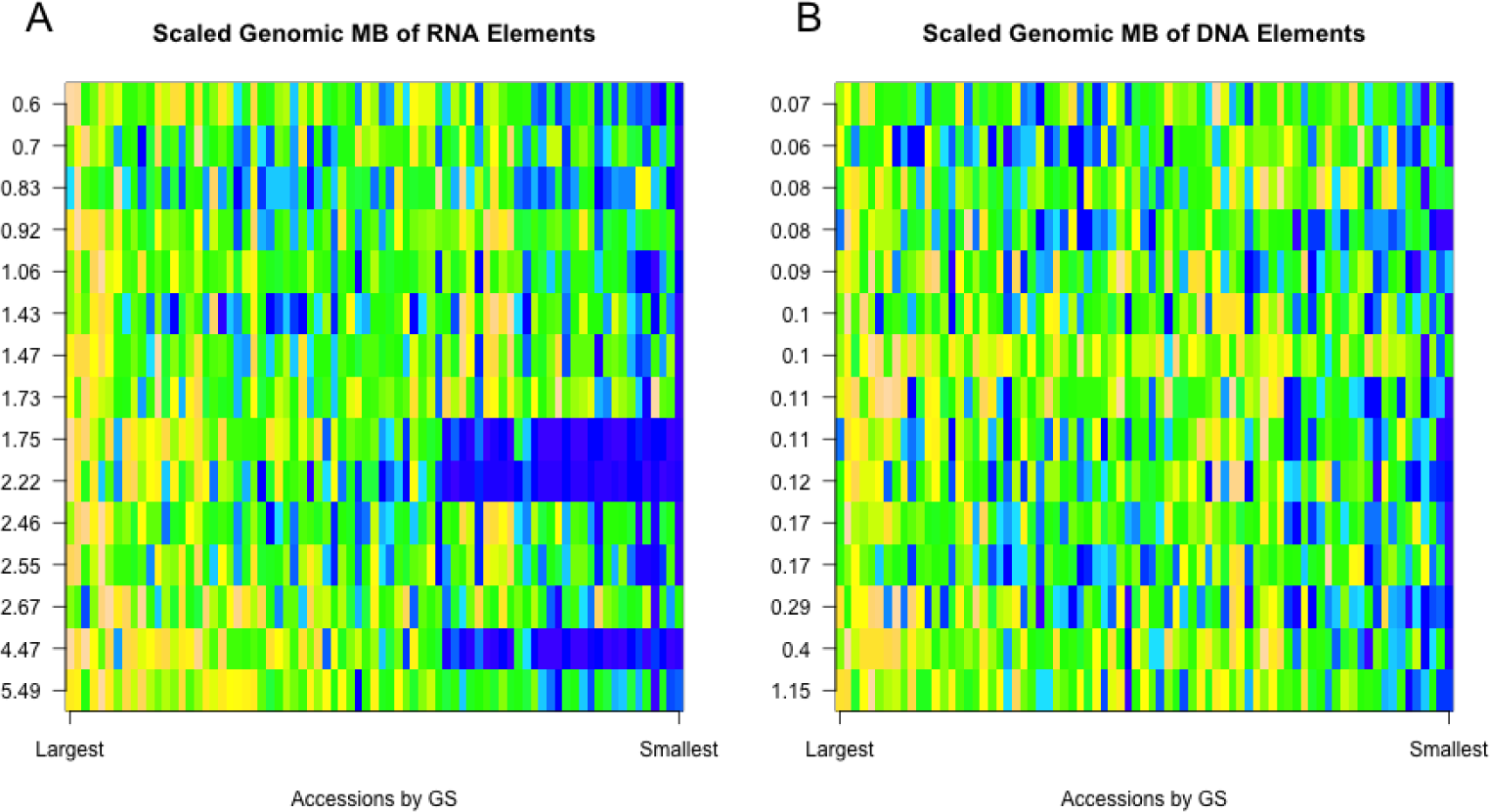
Variation in (A) RNA and (B) DNA transposable element abundance in maize landraces. The y-axis indicates the average abundance in Mb of a given TE subfamily. The fifteen highest abundance subfamilies are shown. The x-axis are maize landraces accessions ordered by genome size, with the largest genome size accessions on the left. Values plotted are bp measures scaled from 0 (blue) to 1 (yellow) per row.

**S6 Fig.**
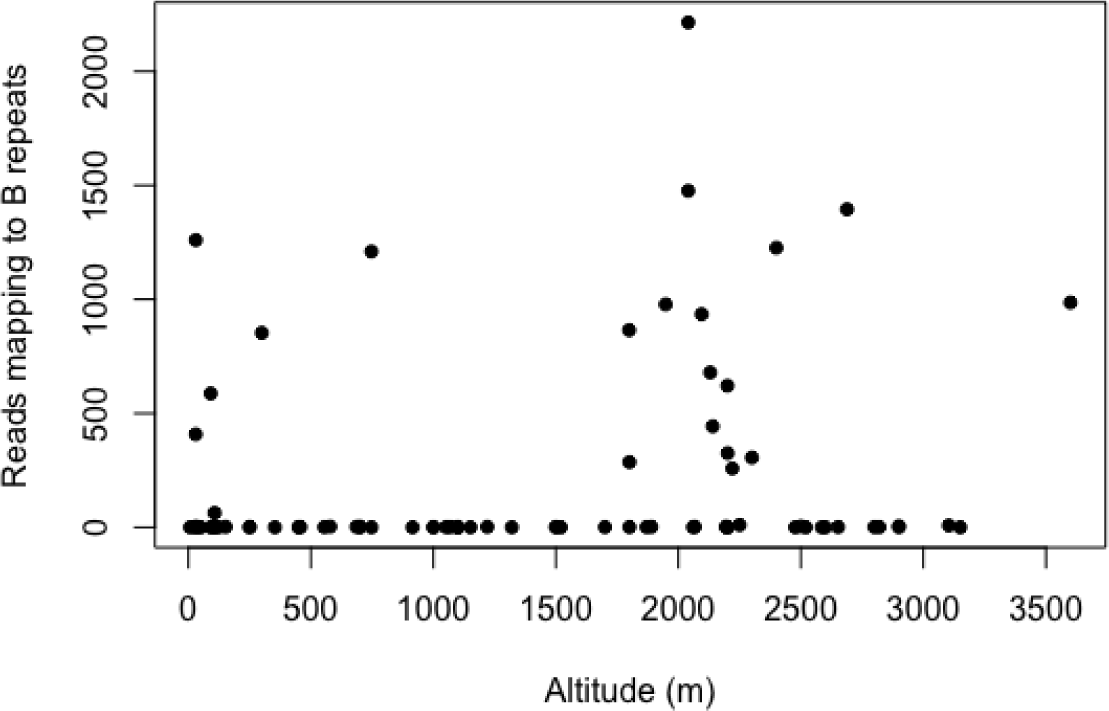
Plot of reads mapping to B chromosome specific repeats in maize landraces.

**S7 Fig.**
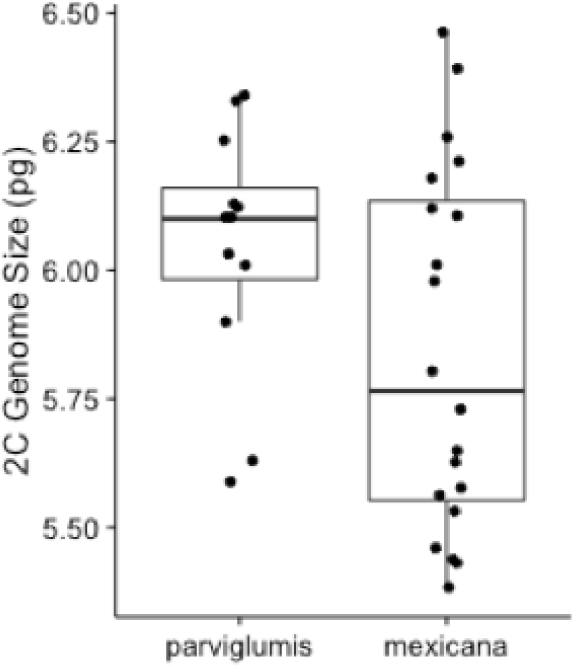
Genome size for two individuals per teosinte population sampled in [37, 50]. Points indicating individual genome size estimates are jittered around the center.

**S8 Fig.**
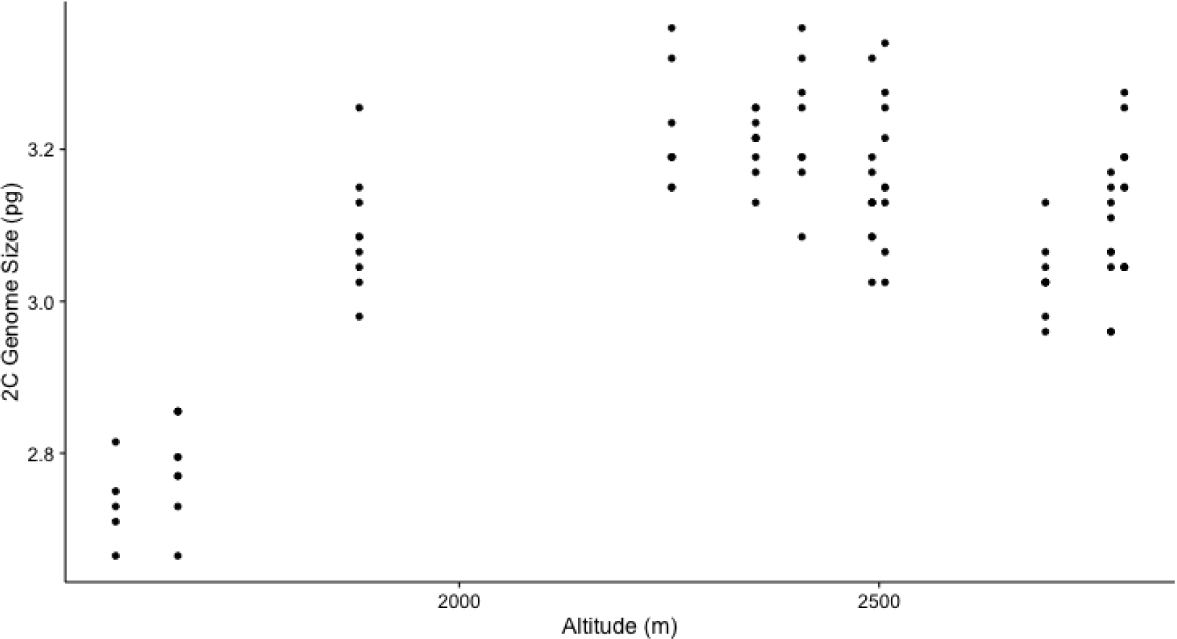
Genome size by altitude of *mexicana*. All samples, including low altitude populations, are shown.

**S9 Fig.**
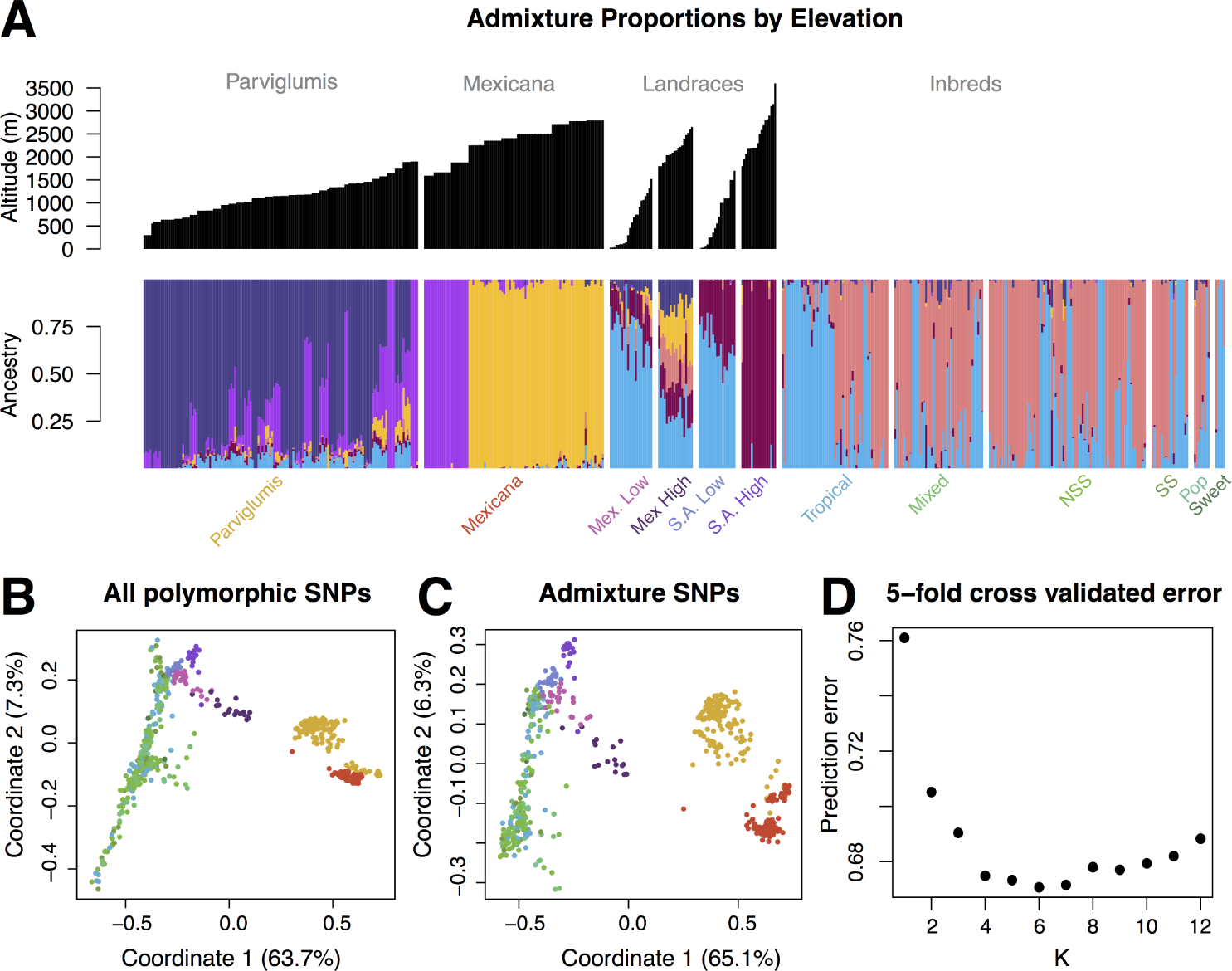
Population structure of maize and *mexicana* populations. (A) Admixture plots for K=6, with altitude of accessions shown above. *Mexicana* populations and maize landraces are those used in this study. We include *parviglumis* [45] and maize inbreds [44]. (B) and (C) Multi-dimensional scaling analyses showing clustering of whole genome SNPs and those used to generate the admixture plot. Points are color coded based on the label underneath the admixture plot. (D) 5-fold cross validated error as estimated by Admixture, indicating the best estimate of number of populations, K.

### S1 Appendix. Path Model

As shown below, a path model enables the relationship between cell production rate and genome size to be inferred by fitting regressions for log CS on log GS, and log LER on log GS. The general approach is described as a model with two “mediators” in Mackinnon [91]. We took a computational Bayesian approach to fitting the path model after early experiments with likelihood-based methods indicated numerical instabilities.

In detail, the first of the regression models is, for cell *c* of seedling *s*:

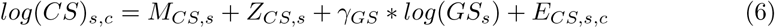

where *M_CS,s_* is a random intercept for the maternal parent of seedling *s*, *Z_CS,s_* is a random intercept for seedling *s*, and *E_CS,s,c_* is a cell-level error term. Inclusion of an overall mean, *μ*, in the regression equation along with the random intercepts led to numerical instabilities apparently related to poor model identification. In effect, the random intercepts– when properly parameterized– take the place of an overall mean. We found that informative priors for *M_CS,s_* and *Z_CS,s_* were necessary: for the final model we used Gaussian priors with mean log(0.003) and standard deviations *σ_MCS_* and *σ_ZCS_*, respectively. This prior mean is the natural logarithm of a typical stomatal cell size, 0.003cm. We expect one of the two random intercepts to assume a greater role in capturing the overall mean. Centering both priors at log(0.003) reflects our indifference to the outcome of this contest. The prior for *E_CS,s,c_* is Gaussian with mean zero and standard deviation *σ_ECS_*. The coefficient *γ_GS_* has a Gaussian prior with mean zero and standard deviation 5.0. We used half-Cauchy priors for the standard deviations *σ_ECS_*, *σ_MCS_* and *σ_ZCS_*.

The second of two regression models– for log LER on log GS– is derived from a model reflecting primary observations of leaf *length* (LL) on successive days of seedling growth. The observation-level model for seedling *s* at time *t* is:

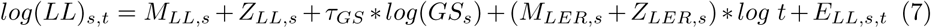

where *M_LL,s_* and *Z_LL,s_* are respectively maternal and seedling random intercepts, *M_LER,s_* and *Z_LER,s_* are maternal and seedling random slopes and *E_LL,s,t_* is an error term. The random slopes *M_LER,s_* and *Z_LER,s_* allow for idiosyncratic growth rates. Natural logarithms on the right- and left-hand sides imply power-law relationships between leaf length, time and genome size in their original units of measurement. As in model 6, informative priors were necessary for model identification. The final model has Gaussian priors with mean log(4.8) and standard deviations *σ_MLL_* and *σ_ZLL_* for *M_LL,s_* and *Z_LL,s_*, respectively. This prior mean is the natural logarithm of a typical leaf elongation rate, 4.8cm– the increment of leaf length that could be expected after a day’s growth (*t* = 1). The priors for *M_LER,s_* and *Z_LER,s_* are Gamma with shape and rate both equal to 1.0 (*i.e.* with mean 1.0, reflecting linear growth). As for model 6, the prior for *E_LL,s,t_* is Gaussian with mean zero and standard deviation *σ_ELL_*, the coefficient *τ_GS_* has a Gaussian prior with mean zero and standard deviation 5.0, and *σ_MLL_*, *σ_ZLL_* and *σ_ELL_* have half-Cauchy priors.

The model for leaf *elongation rate* is subsequently obtained by differentiation of *LL_s,t_* with respect to time:

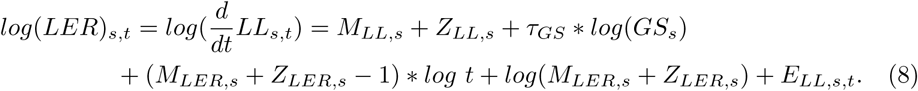

Equations 6 and 8 are placed in context by use of the path diagrams in S2 Fig. Equation 6 is the sub-model connecting log CS to log GS in the left-hand diagram, while equation 8 is a marginal model connecting log LER to log GS, illustrated by the right-hand diagram. The two path diagrams imply two expressions for the same quantity, log LER, which can be equated to produce an estimate of *β_GS_*. The derivation is simplified by taking expected values of the random effects in 6 and 8, and fixing the time horizon at a single day, though the numerical estimates we report come from 6 and 8 in full detail, as displayed above. Subsequently we define

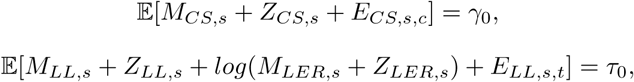

and set *t* = 1 in equation 8. Equations 6 and 8 then simplify as

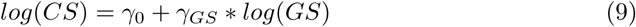

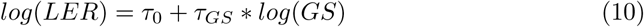

respectively. These are joined by a similar equation for the unobserved variable:

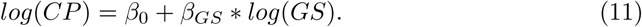

Working from the relationship *LER* = *CS * CP*, or alternatively from log LER backward to its precedents in the left-hand path diagram, we find:

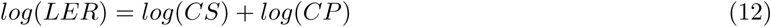

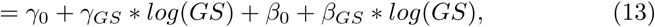

substituting right-hand sides of 9 and 11. Equating expressions 13 and 10 and collecting terms, the coefficient *β_GS_* is recovered as *β_GS_* = *τ_GS_* − *γ_GS_*.

**S1 Table.** Description of data sets used in each analysis.

| Data Group | Genome Size | WGS sequencing Individual | Genotyping Individual | FISH performed? | Used in |
| --- | --- | --- | --- | --- | --- |
| 77 Landraces | Per Individual | Same individual as GS | Different individual than GS but within accession | No | Selection study |
| 93 Mexicana individuals from 11 Populations | Per Individual | Same individual as GS | Same individual as GS | No | Selection study |
| 6 Parviglumis Populations | 2 Individuals per population | NA | NA | No | Teosinte pilot study |
| 10 Mexicana populations | 2 Individuals per population | 9-12 Individuals per population | NA | Yes, different individuals than GS | Teosinte pilot study |

**S2 Table.** Geographic information for maize landrace accessions.

| Accession ID | Region | Altitude (m) | Latitude | Longitude |
| --- | --- | --- | --- | --- |
| RIMMA0388.1 | SAL | 1500 | 6.85 | -75.28333333 |
| RIMMA0389.1 | SAL | 7 | 10.38333333 | -74.88333333 |
| RIMMA0390.1 | SAL | 353 | 4.516666667 | -75.63333333 |
| RIMMA0392.1 | SAL | 555 | 1.75 | -75.58333333 |
| RIMMA0393.1 | SAL | 100 | 8.316666667 | -75.15 |
| RIMMA0394.2 | SAL | 1000 | 4.783333333 | -74.68333333 |
| RIMMA0395.2 | SAL | 30 | 8.5 | -77.26666667 |
| RIMMA0396.1 | SAL | 1100 | 2.583333333 | -75.3 |
| RIMMA0397.1 | SAL | 50 | 11.55 | -72.91666667 |
| RIMMA0398.1 | SAL | 27 | 9.433333333 | -75.7 |
| RIMMA0399.1 | SAL | 250 | 10.18333333 | -74.05 |
| RIMMA0403.2 | SAL | 1000 | 1.25 | -77.51666667 |
| RIMMA0404.1 | SAL | 1500 | 7.3 | -72.51666667 |
| RIMMA0406.1 | SAL | 450 | 4.966666667 | -74.9 |
| RIMMA0409.1 | ML | 107 | 15.43333333 | -92.9 |
| RIMMA0416.1 | MH | 2140 | 29.35 | -107.75 |
| RIMMA0417.1 | MH | 2040 | 28.55 | -107.4833333 |
| RIMMA0418.1 | MH | 2040 | 28.56666667 | -107.4833333 |
| RIMMA0421.1 | MH | 2250 | 19.85 | -97.98333333 |
| RIMMA0422.1 | MH | 2200 | 19.1 | -98.3 |
| RIMMA0423.1 | MH | 2095 | 29.21666667 | -108.1333333 |
| RIMMA0424.1 | MH | 2400 | 27.98333333 | -107.5833333 |
| RIMMA0425.1 | MH | 2130 | 26.81666667 | -107.0666667 |
| RIMMA0426.1 | SAH | 2500 | -9.066666667 | -77.81666667 |
| RIMMA0428.1 | SAL | 700 | -9.3 | -76 |
| RIMMA0430.1 | SAH | 2585 | -9.166666667 | -77.73333333 |
| RIMMA0431.1 | SAH | 2900 | -8.65 | -77.08333333 |
| RIMMA0433.1 | ML | 457 | 14.71666667 | -89.5 |
| RIMMA0436.1 | SAH | 2200 | -6.15 | -77.91666667 |
| RIMMA0437.1 | SAH | 2688 | -9.266666667 | -77.63333333 |
| RIMMA0438.1 | SAH | 2820 | -8.7 | -77.38333333 |
| RIMMA0441.1 | ML | 1320 | 19.16666667 | -96.96666667 |
| RIMMA0462.1 | SAL | 1700 | 1.6 | -77.15 |
| RIMMA0464.1 | SAH | 1800 | -12.33333333 | -74.7 |
| RIMMA0465.1 | SAH | 2300 | -5.566666667 | -79.53333333 |
| RIMMA0466.1 | SAH | 3600 | -14.31666667 | -72.91666667 |
| RIMMA0467.1 | SAH | 2800 | -13.58333333 | -72.91666667 |
| RIMMA0468.1 | SAH | 3150 | -9.383333333 | -77.16666667 |
| RIMMA0473.1 | SAH | 3104 | 1.083333333 | -77.61666667 |
| RIMMA0614.1 | MH | 2060 | 19.95 | -103.7666667 |
| RIMMA0615.1 | ML | 152 | 20.13333333 | -97.2 |
| RIMMA0616.1 | MH | 1800 | 20.81666667 | -102.7666667 |
| RIMMA0619.1 | ML | 747 | 18.35 | -99.53333333 |
| RIMMA0620.1 | MH | 1799 | 20.21666667 | -100.8833333 |
| RIMMA0621.1 | MH | 1870 | 21.11666667 | -101.6833333 |
| RIMMA0623.1 | MH | 2520 | 20.03333333 | -103.6833333 |
| RIMMA0625.1 | MH | 2600 | 19 | -97.38333333 |
| RIMMA0628.1 | ML | 300 | 23.31666667 | -99.01666667 |
| RIMMA0630.1 | MH | 2220 | 19.8 | -97.25 |
| RIMMA0657.1 | SAH | 2201 | -17.5 | -65.66666667 |
| RIMMA0658.1 | SAH | 1948 | -21.83333333 | -64.13333333 |
| RIMMA0661.1 | SAH | 2195 | -2.85 | -78.66666667 |
| RIMMA0662.1 | SAH | 2195 | 0 | -78 |
| RIMMA0663.1 | SAH | 2067 | 0.433333333 | -78.2 |
| RIMMA0667.1 | SAH | 2201 | -21.83333333 | -64.13333333 |
| RIMMA0671.1 | MH | 2477 | 14.76666667 | -91.25 |
| RIMMA0674.1 | MH | 2652 | 19.28333333 | -99.66666667 |
| RIMMA0680.1 | MH | 1890 | 20.36666667 | -102.1666667 |
| RIMMA0690.1 | SAL | 250 | 8.316666667 | -73.63333333 |
| RIMMA0691.1 | SAL | 1098 | 6.55 | -73.13333333 |
| RIMMA0696.1 | ML | 30 | 16.51666667 | -90.16666667 |
| RIMMA0700.1 | ML | 579 | 16.75 | -93.16666667 |
| RIMMA0701.1 | ML | 686 | 16.6 | -92.71666667 |
| RIMMA0702.1 | ML | 1052 | 14.46666667 | -90.75 |
| RIMMA0703.1 | ML | 30 | 20.83333333 | -88.51666667 |
| RIMMA0708.1 | SAL | 1098 | -3.5 | -78.6 |
| RIMMA0709.1 | ML | 747 | 16.5 | -92.5 |
| RIMMA0710.1 | ML | 91 | 15.33333333 | -92.63333333 |
| RIMMA0712.1 | ML | 1220 | 15.28333333 | -90.25 |
| RIMMA0716.1 | ML | 91 | 15.31666667 | -92.66666667 |
| RIMMA0720.1 | ML | 39 | 15.46666667 | -88.85 |
| RIMMA0721.1 | ML | 915 | 14.61666667 | -90.08333333 |
| RIMMA0727.1 | ML | 1151 | 14.4 | -90.46666667 |
| RIMMA0729.1 | ML | 122 | 15.4 | -89.66666667 |
| RIMMA0730.1 | ML | 1067 | 14.48333333 | -90.8 |
| RIMMA0731.1 | ML | 1520 | 16.78333333 | -96.66666667 |
| RIMMA0733.1 | ML | 107 | 16.56666667 | -94.61666667 |

**S3 Table.** Measures of genome size from two individuals from each of the 10 populations used in FISH to sequence correlation (Fig. 2).

| Accession | Subspecies | Ind1 | Ind2 | Altitude(m) |
| --- | --- | --- | --- | --- |
| RIMME0021 | mexicana | 6.12 | 6.01 | 2094 |
| RIMME0026 | mexicana | 6.21 | 6.11 | 2214 |
| RIMME0028 | mexicana | 5.65 | 5.53 | 1916 |
| RIMME0029 | mexicana | 5.46 | 5.58 | 1547 |
| RIMME0030 | mexicana | 5.98 | 5.8 | 2458 |
| RIMME0031 | mexicana | 6.26 | 6.39 | 2609 |
| RIMME0032 | mexicana | 5.63 | 5.73 | 2016 |
| RIMME0033 | mexicana | 5.43 | 5.44 | 1657 |
| RIMME0034 | mexicana | 5.56 | 5.38 | 2173 |
| RIMME0035 | mexicana | 6.18 | 6.46 | 2237 |
| RIMPA0071 | parviglumis | 6.1 | 6.1 | 985 |
| RIMPA0086 | parviglumis | 6.01 | 5.9 | 982 |
| RIMPA0087 | parviglumis | 5.59 | 5.63 | 590 |
| RIMPA0096 | parviglumis | 6.12 | 6.03 | 1528 |
| RIMPA0135 | parviglumis | 6.25 | 6.34 | 880 |
| RIMPA0142 | parviglumis | 6.33 | 6.13 | 1103 |

**S4 Table.**
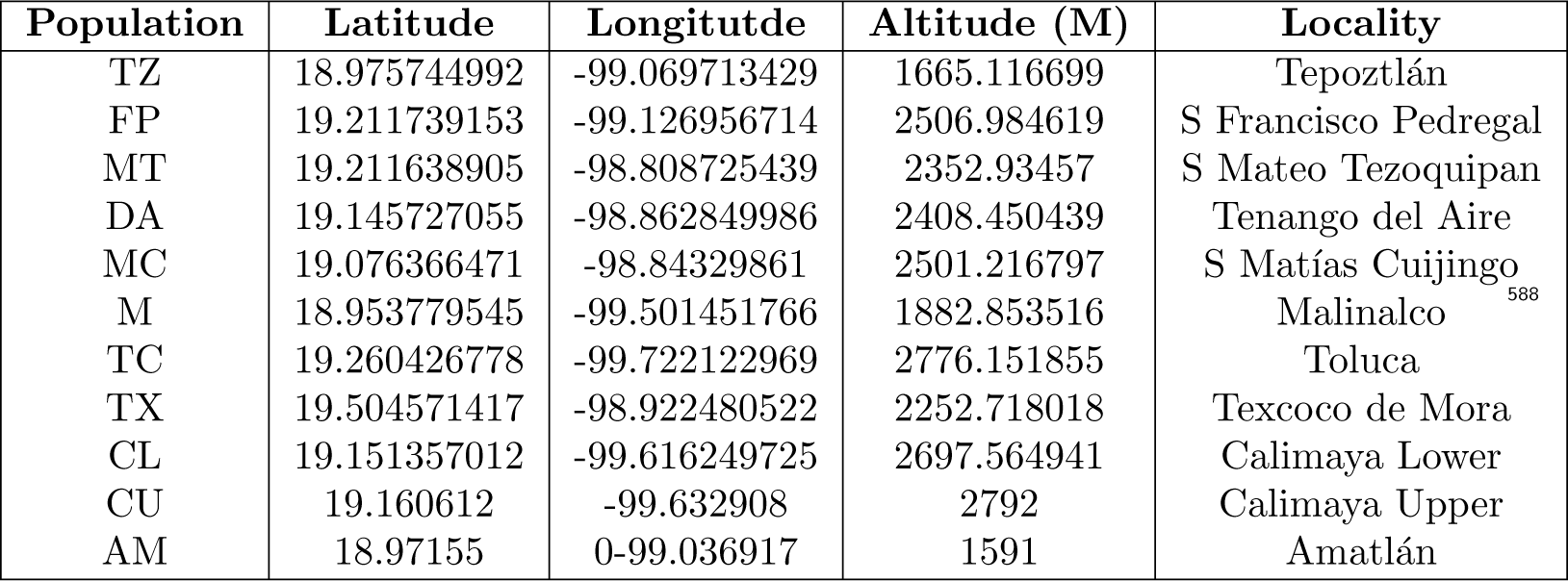
Geographic information for teosinte populations used in selection studies.

**S5 Table.**
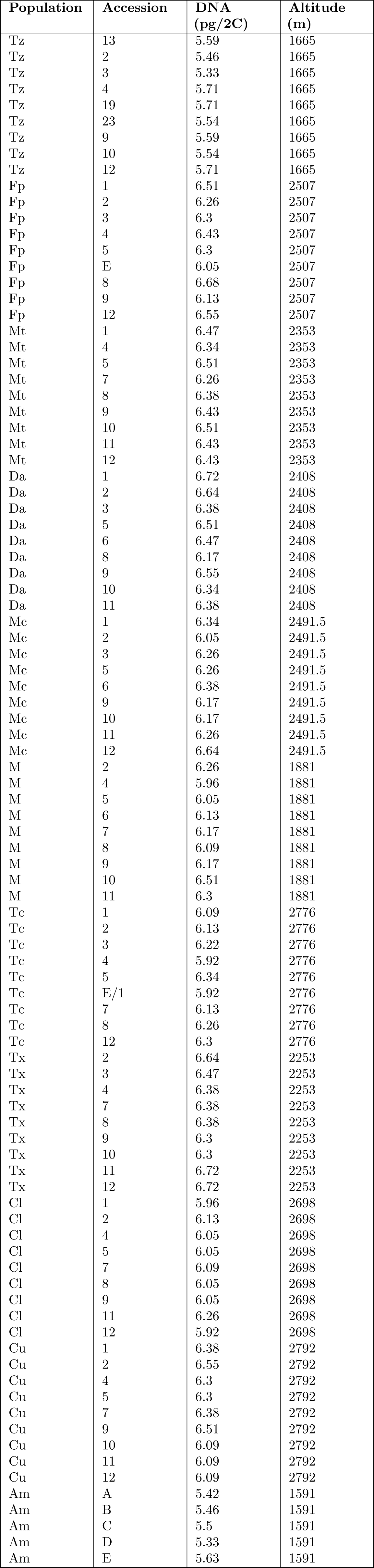
Genome size estimates and altitudinal information for mexicana populations

| Population | Accession | DNA (pg/2C) | Altitude (m) |
| --- | --- | --- | --- |
| Tz | 13 | 5.59 | 1665 |
| Tz | 2 | 5.46 | 1665 |
| Tz | 3 | 5.33 | 1665 |
| Tz | 4 | 5.71 | 1665 |
| Tz | 19 | 5.71 | 1665 |
| Tz | 23 | 5.54 | 1665 |
| Tz | 9 | 5.59 | 1665 |
| Tz | 10 | 5.54 | 1665 |
| Tz | 12 | 5.71 | 1665 |
| Fp | 1 | 6.51 | 2507 |
| Fp | 2 | 6.26 | 2507 |
| Fp | 3 | 6.3 | 2507 |
| Fp | 4 | 6.43 | 2507 |
| Fp | 5 | 6.3 | 2507 |
| Fp | E | 6.05 | 2507 |
| Fp | 8 | 6.68 | 2507 |
| Fp | 9 | 6.13 | 2507 |
| Fp | 12 | 6.55 | 2507 |
| Mt | 1 | 6.47 | 2353 |
| Mt | 4 | 6.34 | 2353 |
| Mt | 5 | 6.51 | 2353 |
| Mt | 7 | 6.26 | 2353 |
| Mt | 8 | 6.38 | 2353 |
| Mt | 9 | 6.43 | 2353 |
| Mt | 10 | 6.51 | 2353 |
| Mt | 11 | 6.43 | 2353 |
| Mt | 12 | 6.43 | 2353 |
| Da | 1 | 6.72 | 2408 |
| Da | 2 | 6.64 | 2408 |
| Da | 3 | 6.38 | 2408 |
| Da | 5 | 6.51 | 2408 |
| Da | 6 | 6.47 | 2408 |
| Da | 8 | 6.17 | 2408 |
| Da | 9 | 6.55 | 2408 |
| Da | 10 | 6.34 | 2408 |
| Da | 11 | 6.38 | 2408 |
| Mc | 1 | 6.34 | 2491.5 |
| Mc | 2 | 6.05 | 2491.5 |
| Mc | 3 | 6.26 | 2491.5 |
| Mc | 5 | 6.26 | 2491.5 |
| Mc | 6 | 6.38 | 2491.5 |
| Mc | 9 | 6.17 | 2491.5 |
| Mc | 10 | 6.17 | 2491.5 |
| Mc | 11 | 6.26 | 2491.5 |
| Mc | 12 | 6.64 | 2491.5 |
| M | 2 | 6.26 | 1881 |
| M | 4 | 5.96 | 1881 |
| M | 5 | 6.05 | 1881 |
| M | 6 | 6.13 | 1881 |
| M | 7 | 6.17 | 1881 |
| M | 8 | 6.09 | 1881 |
| M | 9 | 6.17 | 1881 |
| M | 10 | 6.51 | 1881 |
| M | 11 | 6.3 | 1881 |
| Tc | 1 | 6.09 | 2776 |
| Tc | 2 | 6.13 | 2776 |
| Tc | 3 | 6.22 | 2776 |
| Tc | 4 | 5.92 | 2776 |
| Tc | 5 | 6.34 | 2776 |
| Tc | E/1 | 5.92 | 2776 |
| Tc | 7 | 6.13 | 2776 |
| Tc | 8 | 6.26 | 2776 |
| Tc | 12 | 6.3 | 2776 |
| Tx | 2 | 6.64 | 2253 |
| Tx | 3 | 6.47 | 2253 |
| Tx | 4 | 6.38 | 2253 |
| Tx | 7 | 6.38 | 2253 |
| Tx | 8 | 6.38 | 2253 |
| Tx | 9 | 6.3 | 2253 |
| Tx | 10 | 6.3 | 2253 |
| Tx | 11 | 6.72 | 2253 |
| Tx | 12 | 6.72 | 2253 |
| Cl | 1 | 5.96 | 2698 |
| Cl | 2 | 6.13 | 2698 |
| Cl | 4 | 6.05 | 2698 |
| Cl | 5 | 6.05 | 2698 |
| Cl | 7 | 6.09 | 2698 |
| Cl | 8 | 6.05 | 2698 |
| Cl | 9 | 6.05 | 2698 |
| Cl | 11 | 6.26 | 2698 |
| Cl | 12 | 5.92 | 2698 |
| Cu | 1 | 6.38 | 2792 |
| Cu | 2 | 6.55 | 2792 |
| Cu | 4 | 6.3 | 2792 |
| Cu | 5 | 6.3 | 2792 |
| Cu | 7 | 6.38 | 2792 |
| Cu | 9 | 6.51 | 2792 |
| Cu | 10 | 6.09 | 2792 |
| Cu | 11 | 6.09 | 2792 |
| Cu | 12 | 6.09 | 2792 |
| Am | A | 5.42 | 1591 |
| Am | B | 5.46 | 1591 |
| Am | C | 5.5 | 1591 |
| Am | D | 5.33 | 1591 |
| Am | E | 5.63 | 1591 |

**S6 Table.**
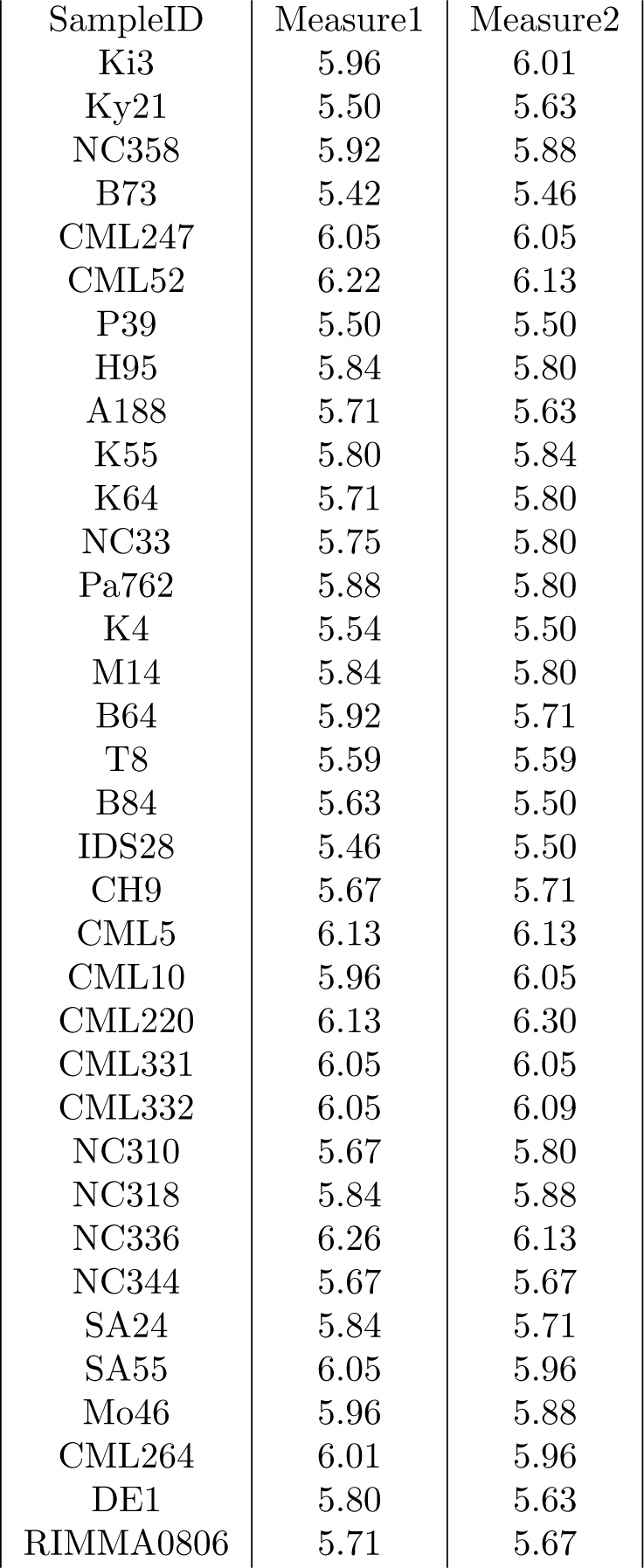
Repeated measures of genome size from maize inbreds lines

| SampleID | Measure1 | Measure2 |
| --- | --- | --- |
| Ki3 | 5.96 | 6.01 |
| Ky21 | 5.50 | 5.63 |
| NC358 | 5.92 | 5.88 |
| B73 | 5.42 | 5.46 |
| CML247 | 6.05 | 6.05 |
| CML52 | 6.22 | 6.13 |
| P39 | 5.50 | 5.50 |
| H95 | 5.84 | 5.80 |
| A188 | 5.71 | 5.63 |
| K55 | 5.80 | 5.84 |
| K64 | 5.71 | 5.80 |
| NC33 | 5.75 | 5.80 |
| Pa762 | 5.88 | 5.80 |
| K4 | 5.54 | 5.50 |
| M14 | 5.84 | 5.80 |
| B64 | 5.92 | 5.71 |
| T8 | 5.59 | 5.59 |
| B84 | 5.63 | 5.50 |
| IDS28 | 5.46 | 5.50 |
| CH9 | 5.67 | 5.71 |
| CML5 | 6.13 | 6.13 |
| CML10 | 5.96 | 6.05 |
| CML220 | 6.13 | 6.30 |
| CML331 | 6.05 | 6.05 |
| CML332 | 6.05 | 6.09 |
| NC310 | 5.67 | 5.80 |
| NC318 | 5.84 | 5.88 |
| NC336 | 6.26 | 6.13 |
| NC344 | 5.67 | 5.67 |
| SA24 | 5.84 | 5.71 |
| SA55 | 6.05 | 5.96 |
| Mo46 | 5.96 | 5.88 |
| CML264 | 6.01 | 5.96 |
| DE1 | 5.80 | 5.63 |
| RIMMA0806 | 5.71 | 5.67 |

**S7 Table.** *Mexicana* Population IDs and number of individuals used for FISH analyses

| Population ID | Number of individuals |
| --- | --- |
| RIMME0021 | 12 |
| RIMME0026 | 12 |
| RIMME0028 | 12 |
| RIMME0029 | 12 |
| RIMME0030 | 12 |
| RIMME0031 | 12 |
| RIMME0032 | 12 |
| RIMME0033 | 12 |
| RIMME0034 | 9 |
| RIMME0035 | 12 |

**S8 Table.** Altitudinal coefficients from selection models using maize landraces and highland teosinte. Calculated altitudinal coefficients (β) from the models testing for altitudinal selection. β values are given in units of megabases per meter. *=p-value <0.05; **=p-value <0.005

| $\beta$ | Maize in MA | Maize in SA | Mexicana |
| --- | --- | --- | --- |
| Genome size | -0.1082932 | -0.1543104 | -0.2698874 |
| Knob180 | -0.004977329 | -0.04344294 | -0.009973119 |
| TR1 | -0.005650659 | -0.00783801 | -0.009217123 |
| TE | -16797840 | 20189310 | 0.07281335 |
| p-value | Maize in MA | Maize in SA | Mexicana |
| Genome size | 0.002860943 | 0.00001214581 | 0.000153224 |
| Knob180 | 0.6955793 | 0.02764277 | 0.9040046 |
| TR1 | 0.02032799 | 0.01453535 | 0.005528401 |
| TE | 0.05931269 | 0.06856391 | 0.1843363 |

**S9 Table.**
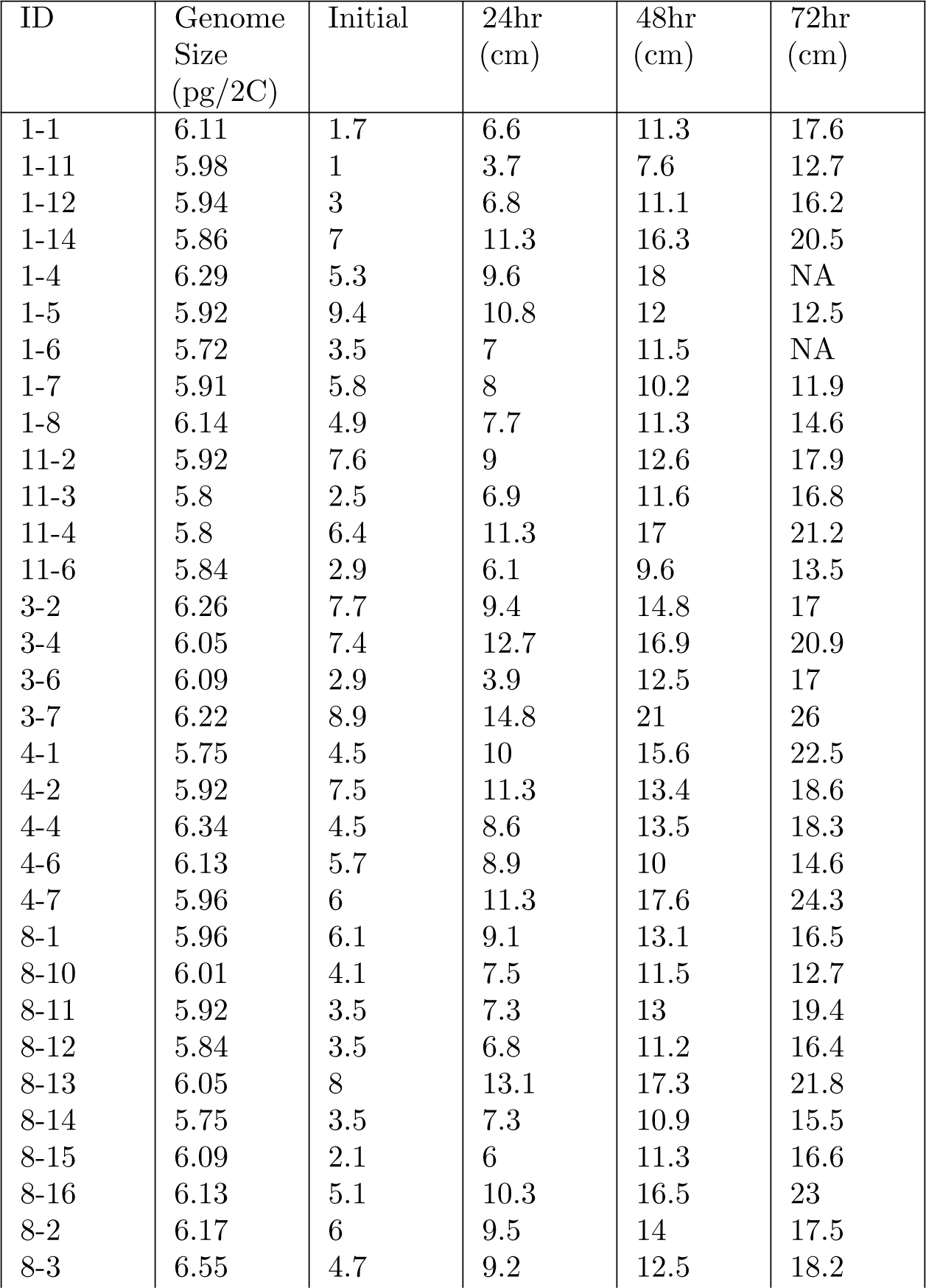
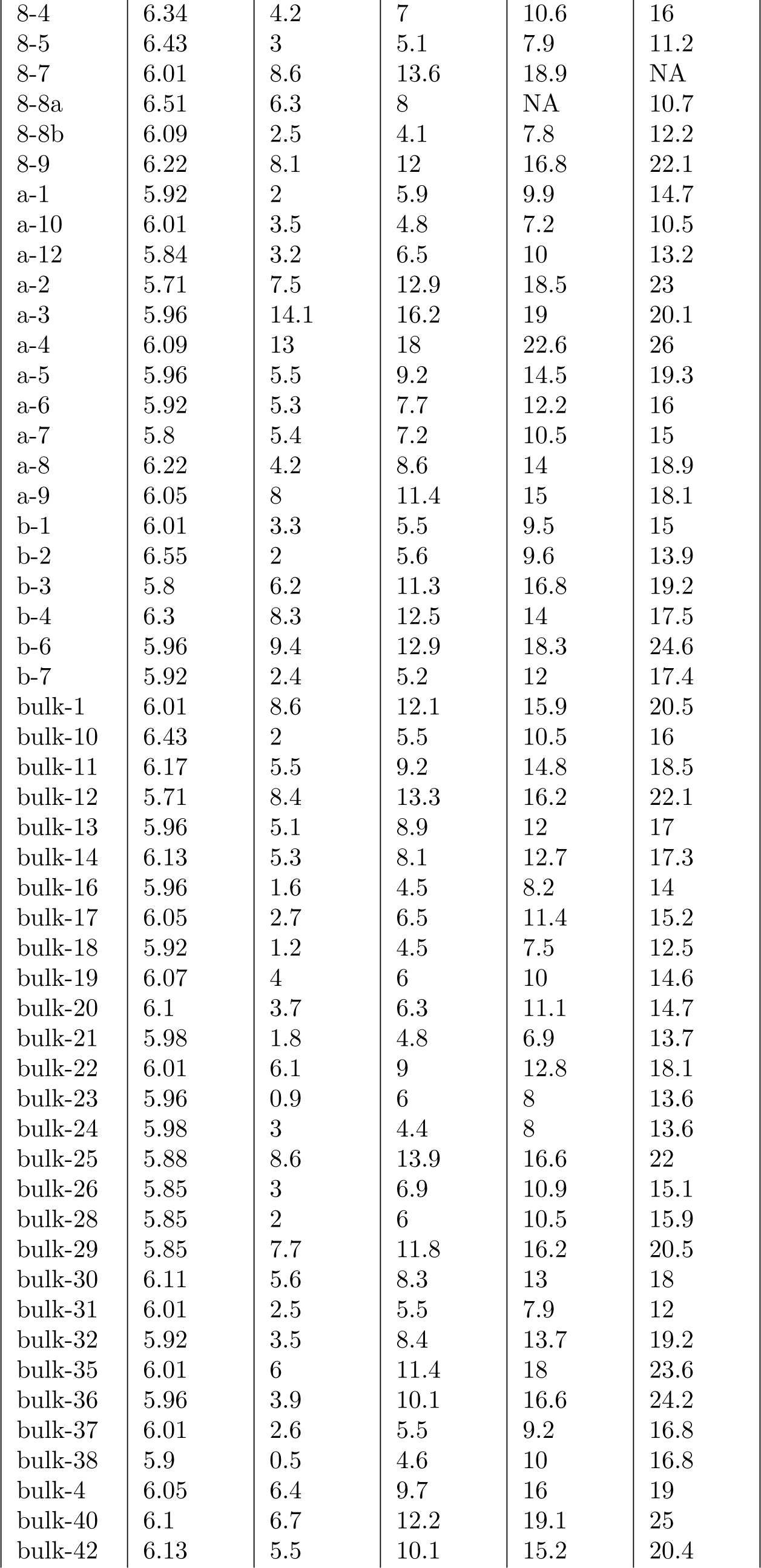
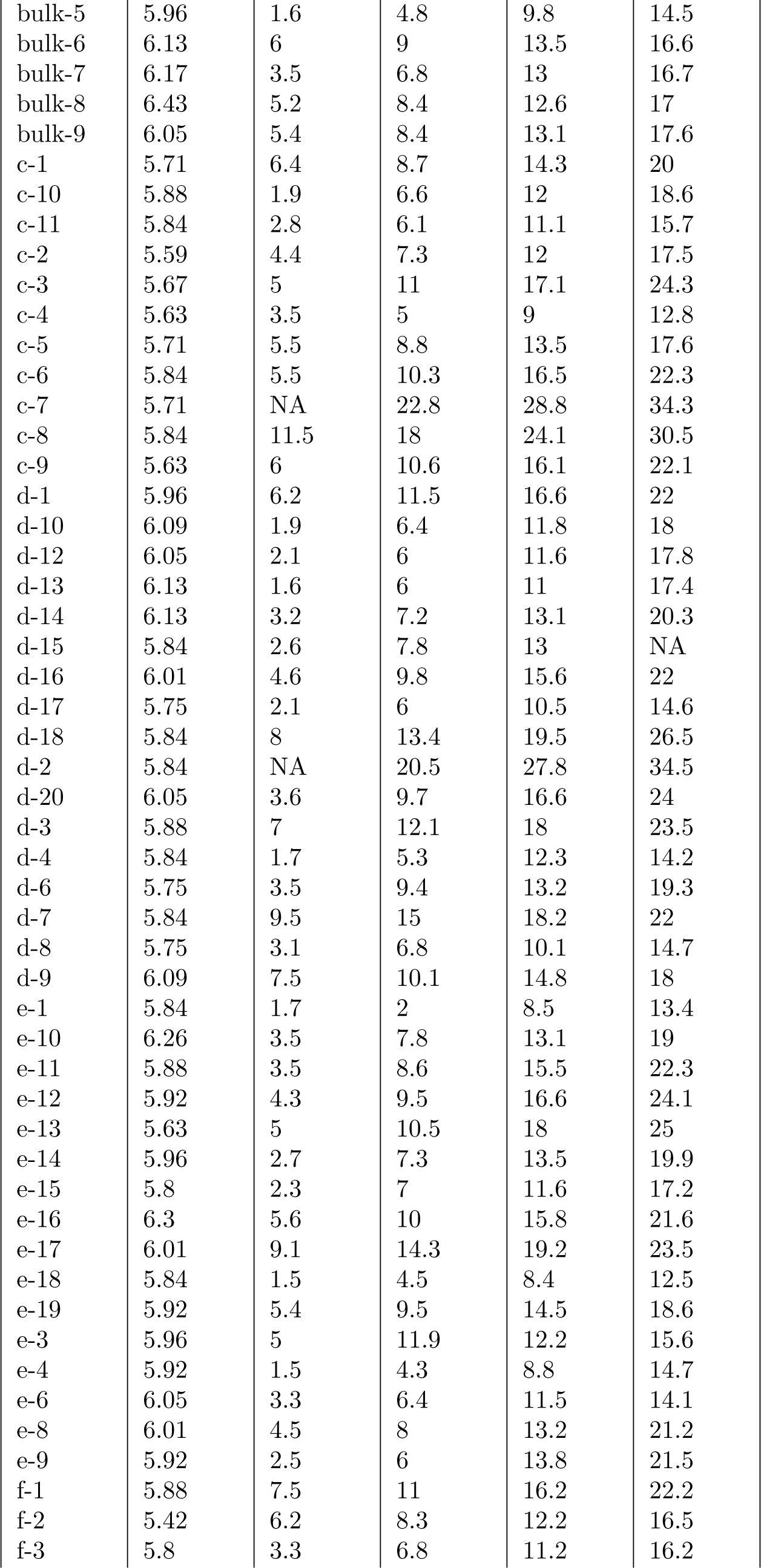
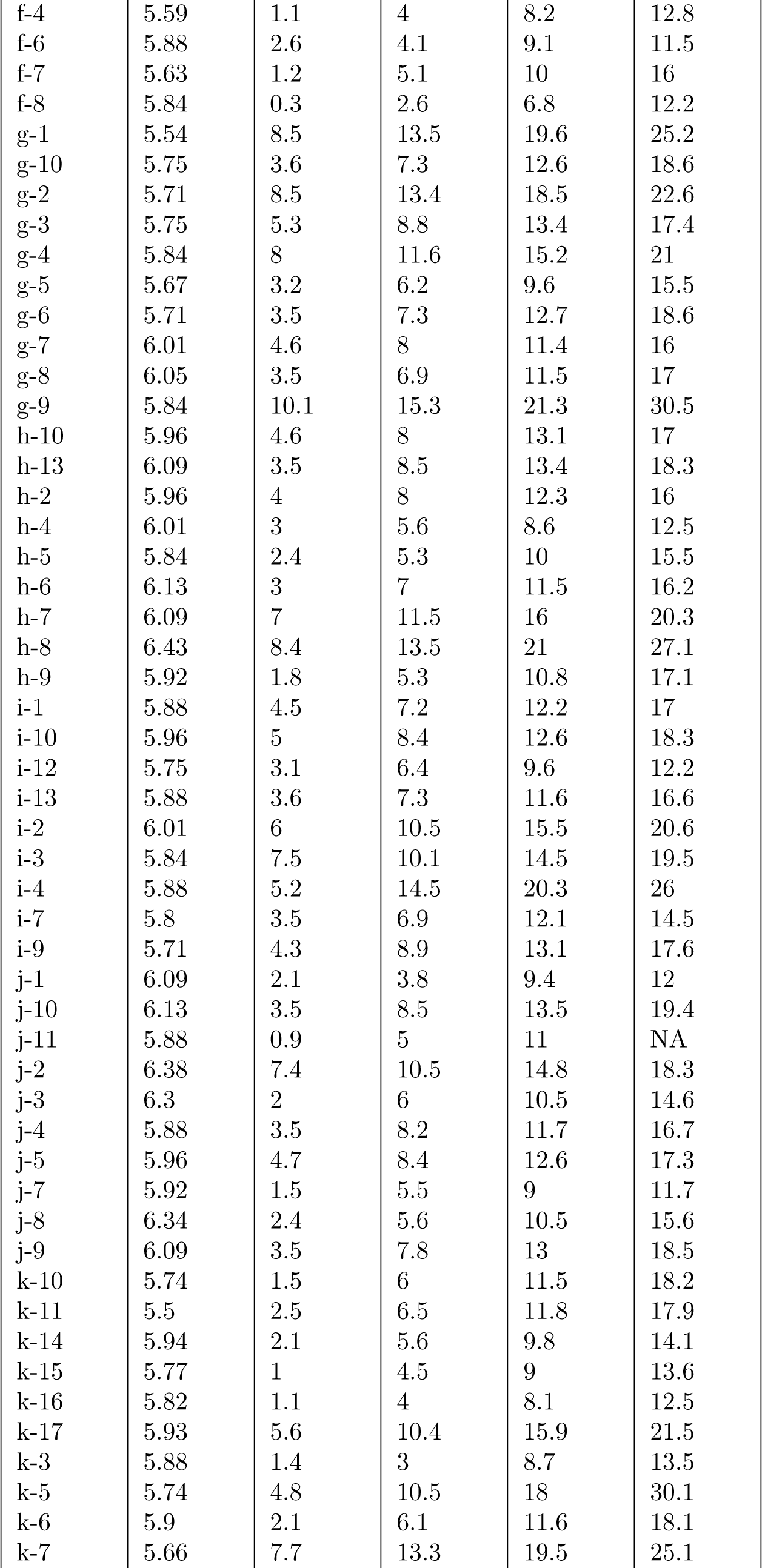
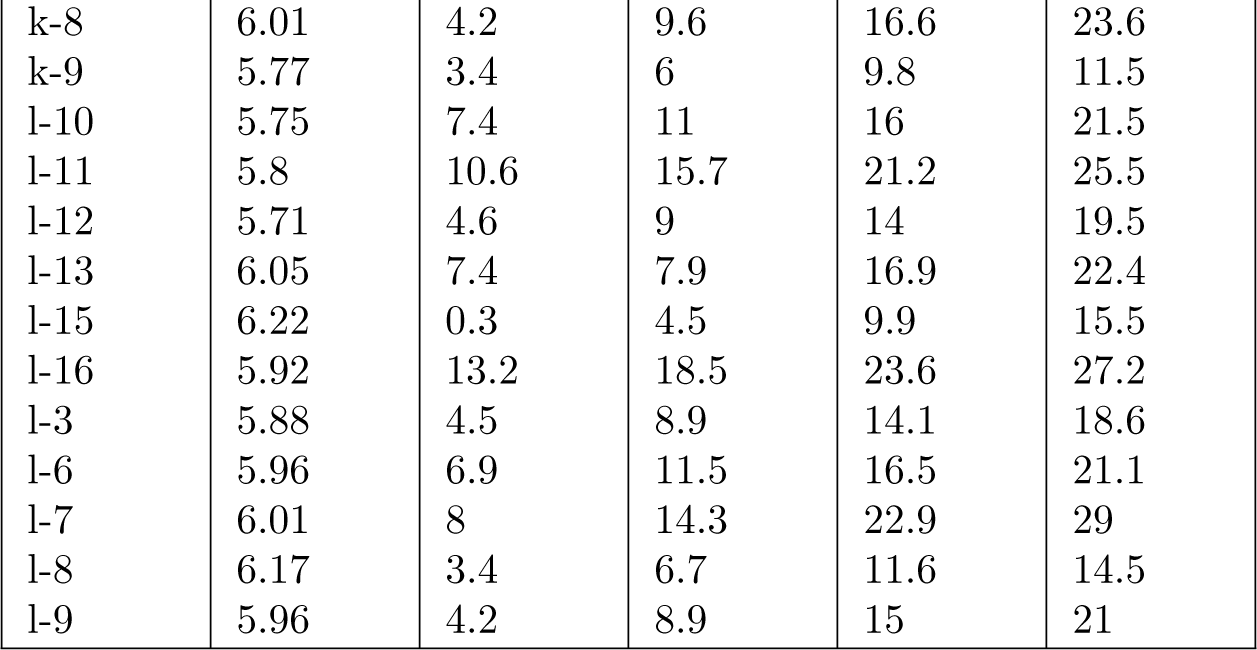
Measured growth rate for Tenango del Aire population in the growth chamber experiment. In plant ID’s, the first digit indicates the mother, while the second is a unique identifier for each individual.

| ID | Genome Size (pg/2C) | Initial | 24hr (cm) | 48hr (cm) | 72hr (cm) |
| --- | --- | --- | --- | --- | --- |
| 1-1 | 6.11 | 1.7 | 6.6 | 11.3 | 17.6 |
| 1-11 | 5.98 | 1 | 3.7 | 7.6 | 12.7 |
| 1-12 | 5.94 | 3 | 6.8 | 11.1 | 16.2 |
| 1-14 | 5.86 | 7 | 11.3 | 16.3 | 20.5 |
| 1-4 | 6.29 | 5.3 | 9.6 | 18 | NA |
| 1-5 | 5.92 | 9.4 | 10.8 | 12 | 12.5 |
| 1-6 | 5.72 | 3.5 | 7 | 11.5 | NA |
| 1-7 | 5.91 | 5.8 | 8 | 10.2 | 11.9 |
| 1-8 | 6.14 | 4.9 | 7.7 | 11.3 | 14.6 |
| 11-2 | 5.92 | 7.6 | 9 | 12.6 | 17.9 |
| 11-3 | 5.8 | 2.5 | 6.9 | 11.6 | 16.8 |
| 11-4 | 5.8 | 6.4 | 11.3 | 17 | 21.2 |
| 11-6 | 5.84 | 2.9 | 6.1 | 9.6 | 13.5 |
| 3-2 | 6.26 | 7.7 | 9.4 | 14.8 | 17 |
| 3-4 | 6.05 | 7.4 | 12.7 | 16.9 | 20.9 |
| 3-6 | 6.09 | 2.9 | 3.9 | 12.5 | 17 |
| 3-7 | 6.22 | 8.9 | 14.8 | 21 | 26 |
| 4-1 | 5.75 | 4.5 | 10 | 15.6 | 22.5 |
| 4-2 | 5.92 | 7.5 | 11.3 | 13.4 | 18.6 |
| 4-4 | 6.34 | 4.5 | 8.6 | 13.5 | 18.3 |
| 4-6 | 6.13 | 5.7 | 8.9 | 10 | 14.6 |
| 4-7 | 5.96 | 6 | 11.3 | 17.6 | 24.3 |
| 8-1 | 5.96 | 6.1 | 9.1 | 13.1 | 16.5 |
| 8-10 | 6.01 | 4.1 | 7.5 | 11.5 | 12.7 |
| 8-11 | 5.92 | 3.5 | 7.3 | 13 | 19.4 |
| 8-12 | 5.84 | 3.5 | 6.8 | 11.2 | 16.4 |
| 8-13 | 6.05 | 8 | 13.1 | 17.3 | 21.8 |
| 8-14 | 5.75 | 3.5 | 7.3 | 10.9 | 15.5 |
| 8-15 | 6.09 | 2.1 | 6 | 11.3 | 16.6 |
| 8-16 | 6.13 | 5.1 | 10.3 | 16.5 | 23 |
| 8-2 | 6.17 | 6 | 9.5 | 14 | 17.5 |
| 8-3 | 6.55 | 4.7 | 9.2 | 12.5 | 18.2 |
| 8-4 | 6.34 | 4.2 | 7 | 10.6 | 16 |
| 8-5 | 6.43 | 3 | 5.1 | 7.9 | 11.2 |
| 8-7 | 6.01 | 8.6 | 13.6 | 18.9 | NA |
| 8-8a | 6.51 | 6.3 | 8 | NA | 10.7 |
| 8-8b | 6.09 | 2.5 | 4.1 | 7.8 | 12.2 |
| 8-9 | 6.22 | 8.1 | 12 | 16.8 | 22.1 |
| a-1 | 5.92 | 2 | 5.9 | 9.9 | 14.7 |
| a-10 | 6.01 | 3.5 | 4.8 | 7.2 | 10.5 |
| a-12 | 5.84 | 3.2 | 6.5 | 10 | 13.2 |
| a-2 | 5.71 | 7.5 | 12.9 | 18.5 | 23 |
| a-3 | 5.96 | 14.1 | 16.2 | 19 | 20.1 |
| a-4 | 6.09 | 13 | 18 | 22.6 | 26 |
| a-5 | 5.96 | 5.5 | 9.2 | 14.5 | 19.3 |
| a-6 | 5.92 | 5.3 | 7.7 | 12.2 | 16 |
| a-7 | 5.8 | 5.4 | 7.2 | 10.5 | 15 |
| a-8 | 6.22 | 4.2 | 8.6 | 14 | 18.9 |
| a-9 | 6.05 | 8 | 11.4 | 15 | 18.1 |
| b-1 | 6.01 | 3.3 | 5.5 | 9.5 | 15 |
| b-2 | 6.55 | 2 | 5.6 | 9.6 | 13.9 |
| b-3 | 5.8 | 6.2 | 11.3 | 16.8 | 19.2 |
| b-4 | 6.3 | 8.3 | 12.5 | 14 | 17.5 |
| b-6 | 5.96 | 9.4 | 12.9 | 18.3 | 24.6 |
| b-7 | 5.92 | 2.4 | 5.2 | 12 | 17.4 |
| bulk-1 | 6.01 | 8.6 | 12.1 | 15.9 | 20.5 |
| bulk-10 | 6.43 | 2 | 5.5 | 10.5 | 16 |
| bulk-11 | 6.17 | 5.5 | 9.2 | 14.8 | 18.5 |
| bulk-12 | 5.71 | 8.4 | 13.3 | 16.2 | 22.1 |
| bulk-13 | 5.96 | 5.1 | 8.9 | 12 | 17 |
| bulk-14 | 6.13 | 5.3 | 8.1 | 12.7 | 17.3 |
| bulk-16 | 5.96 | 1.6 | 4.5 | 8.2 | 14 |
| bulk-17 | 6.05 | 2.7 | 6.5 | 11.4 | 15.2 |
| bulk-18 | 5.92 | 1.2 | 4.5 | 7.5 | 12.5 |
| bulk-19 | 6.07 | 4 | 6 | 10 | 14.6 |
| bulk-20 | 6.1 | 3.7 | 6.3 | 11.1 | 14.7 |
| bulk-21 | 5.98 | 1.8 | 4.8 | 6.9 | 13.7 |
| bulk-22 | 6.01 | 6.1 | 9 | 12.8 | 18.1 |
| bulk-23 | 5.96 | 0.9 | 6 | 8 | 13.6 |
| bulk-24 | 5.98 | 3 | 4.4 | 8 | 13.6 |
| bulk-25 | 5.88 | 8.6 | 13.9 | 16.6 | 22 |
| bulk-26 | 5.85 | 3 | 6.9 | 10.9 | 15.1 |
| bulk-28 | 5.85 | 2 | 6 | 10.5 | 15.9 |
| bulk-29 | 5.85 | 7.7 | 11.8 | 16.2 | 20.5 |
| bulk-30 | 6.11 | 5.6 | 8.3 | 13 | 18 |
| bulk-31 | 6.01 | 2.5 | 5.5 | 7.9 | 12 |
| bulk-32 | 5.92 | 3.5 | 8.4 | 13.7 | 19.2 |
| bulk-35 | 6.01 | 6 | 11.4 | 18 | 23.6 |
| bulk-36 | 5.96 | 3.9 | 10.1 | 16.6 | 24.2 |
| bulk-37 | 6.01 | 2.6 | 5.5 | 9.2 | 16.8 |
| bulk-38 | 5.9 | 0.5 | 4.6 | 10 | 16.8 |
| bulk-4 | 6.05 | 6.4 | 9.7 | 16 | 19 |
| bulk-40 | 6.1 | 6.7 | 12.2 | 19.1 | 25 |
| bulk-42 | 6.13 | 5.5 | 10.1 | 15.2 | 20.4 |
| bulk-5 | 5.96 | 1.6 | 4.8 | 9.8 | 14.5 |
| bulk-6 | 6.13 | 6 | 9 | 13.5 | 16.6 |
| bulk-7 | 6.17 | 3.5 | 6.8 | 13 | 16.7 |
| bulk-8 | 6.43 | 5.2 | 8.4 | 12.6 | 17 |
| bulk-9 | 6.05 | 5.4 | 8.4 | 13.1 | 17.6 |
| c-1 | 5.71 | 6.4 | 8.7 | 14.3 | 20 |
| c-10 | 5.88 | 1.9 | 6.6 | 12 | 18.6 |
| c-11 | 5.84 | 2.8 | 6.1 | 11.1 | 15.7 |
| c-2 | 5.59 | 4.4 | 7.3 | 12 | 17.5 |
| c-3 | 5.67 | 5 | 11 | 17.1 | 24.3 |
| c-4 | 5.63 | 3.5 | 5 | 9 | 12.8 |
| c-5 | 5.71 | 5.5 | 8.8 | 13.5 | 17.6 |
| c-6 | 5.84 | 5.5 | 10.3 | 16.5 | 22.3 |
| c-7 | 5.71 | NA | 22.8 | 28.8 | 34.3 |
| c-8 | 5.84 | 11.5 | 18 | 24.1 | 30.5 |
| c-9 | 5.63 | 6 | 10.6 | 16.1 | 22.1 |
| d-1 | 5.96 | 6.2 | 11.5 | 16.6 | 22 |
| d-10 | 6.09 | 1.9 | 6.4 | 11.8 | 18 |
| d-12 | 6.05 | 2.1 | 6 | 11.6 | 17.8 |
| d-13 | 6.13 | 1.6 | 6 | 11 | 17.4 |
| d-14 | 6.13 | 3.2 | 7.2 | 13.1 | 20.3 |
| d-15 | 5.84 | 2.6 | 7.8 | 13 | NA |
| d-16 | 6.01 | 4.6 | 9.8 | 15.6 | 22 |
| d-17 | 5.75 | 2.1 | 6 | 10.5 | 14.6 |
| d-18 | 5.84 | 8 | 13.4 | 19.5 | 26.5 |
| d-2 | 5.84 | NA | 20.5 | 27.8 | 34.5 |
| d-20 | 6.05 | 3.6 | 9.7 | 16.6 | 24 |
| d-3 | 5.88 | 7 | 12.1 | 18 | 23.5 |
| d-4 | 5.84 | 1.7 | 5.3 | 12.3 | 14.2 |
| d-6 | 5.75 | 3.5 | 9.4 | 13.2 | 19.3 |
| d-7 | 5.84 | 9.5 | 15 | 18.2 | 22 |
| d-8 | 5.75 | 3.1 | 6.8 | 10.1 | 14.7 |
| d-9 | 6.09 | 7.5 | 10.1 | 14.8 | 18 |
| e-1 | 5.84 | 1.7 | 2 | 8.5 | 13.4 |
| e-10 | 6.26 | 3.5 | 7.8 | 13.1 | 19 |
| e-11 | 5.88 | 3.5 | 8.6 | 15.5 | 22.3 |
| e-12 | 5.92 | 4.3 | 9.5 | 16.6 | 24.1 |
| e-13 | 5.63 | 5 | 10.5 | 18 | 25 |
| e-14 | 5.96 | 2.7 | 7.3 | 13.5 | 19.9 |
| e-15 | 5.8 | 2.3 | 7 | 11.6 | 17.2 |
| e-16 | 6.3 | 5.6 | 10 | 15.8 | 21.6 |
| e-17 | 6.01 | 9.1 | 14.3 | 19.2 | 23.5 |
| e-18 | 5.84 | 1.5 | 4.5 | 8.4 | 12.5 |
| e-19 | 5.92 | 5.4 | 9.5 | 14.5 | 18.6 |
| e-3 | 5.96 | 5 | 11.9 | 12.2 | 15.6 |
| e-4 | 5.92 | 1.5 | 4.3 | 8.8 | 14.7 |
| e-6 | 6.05 | 3.3 | 6.4 | 11.5 | 14.1 |
| e-8 | 6.01 | 4.5 | 8 | 13.2 | 21.2 |
| e-9 | 5.92 | 2.5 | 6 | 13.8 | 21.5 |
| f-1 | 5.88 | 7.5 | 11 | 16.2 | 22.2 |
| f-2 | 5.42 | 6.2 | 8.3 | 12.2 | 16.5 |
| f-3 | 5.8 | 3.3 | 6.8 | 11.2 | 16.2 |
| f-4 | 5.59 | 1.1 | 4 | 8.2 | 12.8 |
| f-6 | 5.88 | 2.6 | 4.1 | 9.1 | 11.5 |
| f-7 | 5.63 | 1.2 | 5.1 | 10 | 16 |
| f-8 | 5.84 | 0.3 | 2.6 | 6.8 | 12.2 |
| g-1 | 5.54 | 8.5 | 13.5 | 19.6 | 25.2 |
| g-10 | 5.75 | 3.6 | 7.3 | 12.6 | 18.6 |
| g-2 | 5.71 | 8.5 | 13.4 | 18.5 | 22.6 |
| g-3 | 5.75 | 5.3 | 8.8 | 13.4 | 17.4 |
| g-4 | 5.84 | 8 | 11.6 | 15.2 | 21 |
| g-5 | 5.67 | 3.2 | 6.2 | 9.6 | 15.5 |
| g-6 | 5.71 | 3.5 | 7.3 | 12.7 | 18.6 |
| g-7 | 6.01 | 4.6 | 8 | 11.4 | 16 |
| g-8 | 6.05 | 3.5 | 6.9 | 11.5 | 17 |
| g-9 | 5.84 | 10.1 | 15.3 | 21.3 | 30.5 |
| h-10 | 5.96 | 4.6 | 8 | 13.1 | 17 |
| h-13 | 6.09 | 3.5 | 8.5 | 13.4 | 18.3 |
| h-2 | 5.96 | 4 | 8 | 12.3 | 16 |
| h-4 | 6.01 | 3 | 5.6 | 8.6 | 12.5 |
| h-5 | 5.84 | 2.4 | 5.3 | 10 | 15.5 |
| h-6 | 6.13 | 3 | 7 | 11.5 | 16.2 |
| h-7 | 6.09 | 7 | 11.5 | 16 | 20.3 |
| h-8 | 6.43 | 8.4 | 13.5 | 21 | 27.1 |
| h-9 | 5.92 | 1.8 | 5.3 | 10.8 | 17.1 |
| i-1 | 5.88 | 4.5 | 7.2 | 12.2 | 17 |
| i-10 | 5.96 | 5 | 8.4 | 12.6 | 18.3 |
| i-12 | 5.75 | 3.1 | 6.4 | 9.6 | 12.2 |
| i-13 | 5.88 | 3.6 | 7.3 | 11.6 | 16.6 |
| i-2 | 6.01 | 6 | 10.5 | 15.5 | 20.6 |
| i-3 | 5.84 | 7.5 | 10.1 | 14.5 | 19.5 |
| i-4 | 5.88 | 5.2 | 14.5 | 20.3 | 26 |
| i-7 | 5.8 | 3.5 | 6.9 | 12.1 | 14.5 |
| i-9 | 5.71 | 4.3 | 8.9 | 13.1 | 17.6 |
| j-1 | 6.09 | 2.1 | 3.8 | 9.4 | 12 |
| j-10 | 6.13 | 3.5 | 8.5 | 13.5 | 19.4 |
| j-11 | 5.88 | 0.9 | 5 | 11 | NA |
| j-2 | 6.38 | 7.4 | 10.5 | 14.8 | 18.3 |
| j-3 | 6.3 | 2 | 6 | 10.5 | 14.6 |
| j-4 | 5.88 | 3.5 | 8.2 | 11.7 | 16.7 |
| j-5 | 5.96 | 4.7 | 8.4 | 12.6 | 17.3 |
| j-7 | 5.92 | 1.5 | 5.5 | 9 | 11.7 |
| j-8 | 6.34 | 2.4 | 5.6 | 10.5 | 15.6 |
| j-9 | 6.09 | 3.5 | 7.8 | 13 | 18.5 |
| k-10 | 5.74 | 1.5 | 6 | 11.5 | 18.2 |
| k-11 | 5.5 | 2.5 | 6.5 | 11.8 | 17.9 |
| k-14 | 5.94 | 2.1 | 5.6 | 9.8 | 14.1 |
| k-15 | 5.77 | 1 | 4.5 | 9 | 13.6 |
| k-16 | 5.82 | 1.1 | 4 | 8.1 | 12.5 |
| k-17 | 5.93 | 5.6 | 10.4 | 15.9 | 21.5 |
| k-3 | 5.88 | 1.4 | 3 | 8.7 | 13.5 |
| k-5 | 5.74 | 4.8 | 10.5 | 18 | 30.1 |
| k-6 | 5.9 | 2.1 | 6.1 | 11.6 | 18.1 |
| k-7 | 5.66 | 7.7 | 13.3 | 19.5 | 25.1 |
| k-8 | 6.01 | 4.2 | 9.6 | 16.6 | 23.6 |
| k-9 | 5.77 | 3.4 | 6 | 9.8 | 11.5 |
| l-10 | 5.75 | 7.4 | 11 | 16 | 21.5 |
| l-11 | 5.8 | 10.6 | 15.7 | 21.2 | 25.5 |
| l-12 | 5.71 | 4.6 | 9 | 14 | 19.5 |
| l-13 | 6.05 | 7.4 | 7.9 | 16.9 | 22.4 |
| l-15 | 6.22 | 0.3 | 4.5 | 9.9 | 15.5 |
| l-16 | 5.92 | 13.2 | 18.5 | 23.6 | 27.2 |
| l-3 | 5.88 | 4.5 | 8.9 | 14.1 | 18.6 |
| l-6 | 5.96 | 6.9 | 11.5 | 16.5 | 21.1 |
| l-7 | 6.01 | 8 | 14.3 | 22.9 | 29 |
| l-8 | 6.17 | 3.4 | 6.7 | 11.6 | 14.5 |
| l-9 | 5.96 | 4.2 | 8.9 | 15 | 21 |

## Acknowledgments

We would like to thank Arvid Ågren, Graham Coop, Peter Ralph, Michelle Stitzer, Hernàn Burbano, as well as members of the Ross-Ibarra, Coop, and Burbano labs for helpful discussion. We thank Anna O’Brien for providing seed from her *mexicana* collections and for early access to her SNP data. We acknowledge financial support from NSF grants IOS-0922703 and IOS-1238014 and the USDA Hatch project CA-D-PLS-2066-H. P.B. would like to thank the UC MEXUS Dissertation Grant, DuPont Pioneer, and the UC Davis Department of Plant Sciences for funding and support.

